# Using distance on the Riemannian manifold to compare representations in brain and models

**DOI:** 10.1101/2020.11.25.398511

**Authors:** Mahdiyar Shahbazi, Ali Shirali, Hamid Aghajan, Hamed Nili

**Author notes:** Joint first authors (Mahdiyar Shahbazi, Ali Shirali).

## Abstract

Representational similarity analysis (RSA) summarizes activity patterns for a set of experimental conditions into a matrix composed of pairwise comparisons between activity patterns. Two examples of such matrices are the condition-by-condition inner product matrix or the correlation matrix. These representational matrices reside on the manifold of positive semidefinite matrices, called the Riemannian manifold. We hypothesize that representational similarities would be more accurately quantified by considering the underlying manifold of the representational matrices. Thus, we introduce the distance on the Riemannian manifold as a metric for comparing representations. Analyzing simulated and real fMRI data and considering a wide range of metrics, we show that the Riemannian distance is least susceptible to sampling bias, results in larger intra-subject reliability, and affords searchlight mapping with high sensitivity and specificity. Furthermore, we show that the Riemannian distance can be used for measuring multi-dimensional connectivity. This measure captures both univariate and multivariate connectivity and is also more sensitive to nonlinear regional interactions compared to the state-of-the-art measures. Applying our proposed metric to neural network representations of natural images, we demonstrate that it also possesses outstanding performance in quantifying similarity in models. Taken together, our results lend credence to the proposition that RSA should consider the manifold of the representational matrices to summarize response patterns in the brain and models.

## 1 Introduction

Investigating the information content of brain representations helps in understanding the functional role of different brain areas. Pattern-information analysis has benefitted from a number of analysis techniques that lie at the intersection of neuroscience and machine learning. Generally, these techniques can be grouped into two main categories: pattern classifiers and representational methods.

In pattern classifiers (Haynes and Rees, 2006) successful classification of activity patterns implies information about particular distinctions. In representational methods, i.e. encoding analysis (Kay et al., 2008), representational similarity analysis (RSA; Kriegeskorte et al., 2008) and pattern-component modelling (PCM; Diedrichsen et al., 2011), distributed activity patterns for a number of conditions in a region are summarized by a condition-by-condition inner product matrix, aka the 2^nd^-moment matrix of activity patterns (Diedrichsen and Kriegeskorte, 2017). In particular, RSA exploits representational matrices composed of (dis)similarities between activity patterns for all pairs of conditions. Some examples of theses matrices are representational dissimilarity matrices (RDMs), stimulus-by-stimulus correlation matrices^1^, and also 2^nd^-moment matrices.

2^nd^-moment and cross-correlation matrices, and generally all positive semi-definite (PSD) matrices, lie on a manifold. If equipped with a Riemannian metric, the manifold is called a “Riemannian manifold”. This suggests that the relationship between different PSD matrices would be best captured if the geometry of the manifold is considered.

Considering the geometry of PSD matrices as a Riemannian geometry dates back to (Pennec, 2006). He proposed an affine-invariant Riemannian metric on the manifold of PSD matrices. Several domains in neuroscience research involve correlation or covariance matrices, which are all PSD matrices thus do not form a Euclidean space in nature. Therefore, researchers have started to revise existing approaches in such areas. Examples of such areas are diffusion tensor imaging (Pennec et al., 2006), brain-computer interface classification (Barachant et al., 2012), functional connectivity (Pervaiz et al., 2020; You and Park, 2021) and covariance shrinkage techniques (Rahim et al., 2019). The proposed metric enjoys a number of properties including being affine-invariant (Pennec et al., 2019) and robust to outliers (Congedo et al., 2017).

As a simple demonstration, we use a toy example to show that the relationship between three PSD matrices would be different depending on whether the underlying geometry of the matrices is considered or not. Let Σ_1_, Σ_2_, and Σ_3_ be three 2 × 2 PSD matrices as illustrated in **Figure 1**. As these matrices are composed of three unique elements, each matrix could be visualized as a point in ℝ^3^.

**Figure 1.**
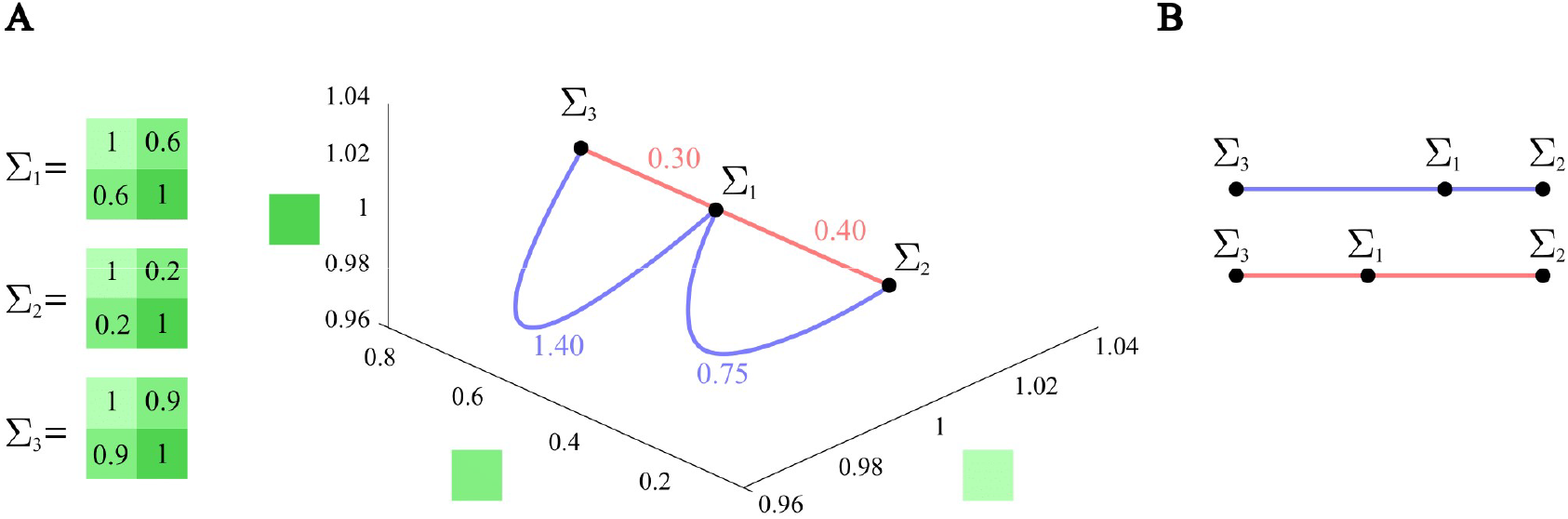
Relationship between PSD matrices can differ when the underlying Riemannian manifold is considered. ***A.** Here, Σ*_1_, *Σ*_2_*, and Σ*_3_ *are three PSD matrices. The numbers written on the blue curves are the lengths of the shortest paths on the manifold of PSD matrices connecting them. It is worth noting that the lengths of blue curves are calculated according to the Riemannian metric, which is different from their lengths in the Euclidean space. In contrast, the numbers written on the red straight lines are the Euclidean distances between matrices (both diagonal and lower-diagonal elements). In this illustration, we only show the diagonal and lower-diagonal elements since the matrices are symmetric. **B.** 1D visualization of relative positions of Σ*_1_, *Σ*_2_*, and Σ*_3_ *using the Riemannian distance (blue) or Euclidean distance (red)*.

The blue curves in **Figure 1** correspond to the shortest paths connecting Σ_1_ to Σ_3_ and Σ_1_ to Σ_2_ on the Riemannian manifold. One can easily see that Σ_1_ is closer to Σ_3_ in the 3-dimensional Euclidean space (0.3 vs. 0.4), while when we consider the embedding manifold of the matrices, Σ_1_ is closer to Σ_2_ (0.75 vs. 1.40).

These kinds of comparisons are frequently done when representational models are being tested (Kriegeskorte and Kievit, 2013). Let it be a brain region or a layer of a computational neural network, considering the geometry can result in different relationships, and this can affect conclusions about representational similarities.

In this paper, we investigate the advantages of considering the underlying geometry of all PSD matrices when quantifying the relationship between representational matrices. To this end, we use simulations with known underlying effects and also fMRI data from two publicly available datasets, Visual object recognition dataset (Hanson et al., 2004; Haxby et al., 2001; O’Toole et al., 2005), and Generic object decoding dataset (Horikawa and Kamitani, 2017).

We first provide a concise introduction to Riemannian geometry and the distance between two PSD matrices. Using simulations and real data, we find domains in which comparing representational matrices via the Riemannian distance significantly improves RSA.

In brief, we will verify the following advantages in testing representational models with the Riemannian metric: this metric allows for more accurate representational comparisons for a relatively small number of response channels. It also yields more consistent within-subject results, thus offering higher reliability. Additionally, representational structures from different brain areas are more discriminable when compared using the Riemannian distance. We also explore the advantages of using the Riemannian metric for representational connectivity (where the aim is to compare the representational content for a set of conditions in distributed activity patterns of two brain regions). We show that our proposed metric goes beyond existing methods (Basti et al., 2020) in capturing nonlinear regional interactions as well as linear and multivariate as well as univariate functional connectivity. We then analyse image representations in different layers of deep neural networks and show that the Riemannian metric can be used to quantify the similarity of neural network representations with higher accuracy than state-of-the-art methods (Kornblith et al., 2019). Finally, we show that when adopted for searchlight mapping, this metric performs well and offers reasonable sensitivity.

## 2 Materials and methods

Throughout the paper, we rely on one of the main points explained in-depth in (Diedrichsen and Kriegeskorte, 2017). Namely, the fact that all methods that enable testing representational models, rely on the 2^nd^-moment matrices. In particular, considering the stimulus-response matrix *U*, which is *k* × *p*, where *k* is the number of experimental conditions and *p* is the number of response channels, the 2^nd^-moment matrix, G, is defined as *UU*^*T*^/*p*. Now, representational dissimilarity matrices (RDMs) could be derived directly from G. Further, representational similarity matrices (RSMs), which are the *k* × *k* correlation matrices, could be derived from G^2^ as well. We have already mentioned that RSMs are closely related to correlation-distance RDMs: RSM = 1 - RDM.RDMs, RSMs, and 2^nd^-moment matrices could be used to characterize multivariate response-patterns of a brain region. RSMs contain the same information as correlation-distance RDMs. However, 2^nd^ moment matrices are more general. In this manuscript we consider both RSMs and 2^nd^-moment matrices. The motivation for including RSMs is to relate to the classic RSA inference.

### 2.1 Metrics for comparing representations

In this section, we investigate a number of metrics for comparing representational structures.

**Table 1** lists the different metrics along with their descriptions and mathematical definitions to calculate the distance or correlation between two *k* × *k* PSD matrices, Σ_1_ and Σ_2_. The correlation-based methods listed in **Table 1** use vectorized matrices. The vectorizing operation (*vec*(.)) converts the lower triangular part of a matrix to a vector, which might include the diagonal or not depending on the type of the matrix (2^nd^-moment matrix or RSM; section 2.3). Let’s call the vectorized Σ_1_ and Σ_2_, **v**_**1**_ and **v**_**2**_, respectively. We show the *i*^th^ element of vector **v**_**1**_ (**v**_**2**_) by *v*_1,*i*_ (*v*_2,*i*_).

**Table 1.** List of all investigated metrics.

| Metric | Description | Formula |
| --- | --- | --- |
| Riemannian distance | The length of the shortest path (geodesic) connecting two PSD matrices on the Riemannian manifold | $\sqrt{\sum_{i=1}^k \log^2(\sigma_i)}$<br>where $\sigma_i$ is the $i^{th}$ eigenvalue of $\Sigma_1^{-1}\Sigma_2$. |
| Pearson correlation | The sample covariance (*cov*) of two vectors normalized by the product of their standard deviations (*std*) | $\frac{cov(\mathbf{v}_1, \mathbf{v}_2)}{std(\mathbf{v}_1) std(\mathbf{v}_2)}$ |
| Spearman correlation | The Pearson correlation between rank-transformed (*rank*) vectors | $\frac{cov(rank(\mathbf{v}_1), rank(\mathbf{v}_2))}{std(rank(\mathbf{v}_1)) std(rank(\mathbf{v}_2))}$ |
| Kendall's $\tau$ correlation | A measure of similarity of orderings of two vectors | $\frac{2}{n(n-1)} \sum_{i=1}^{n-1} \sum_{j=i+1}^n sign(v_{1,i} - v_{1,j}) sign(v_{2,i} - v_{2,j})$<br>$n = k(k+1)/2$ |
| Frobenius norm of the difference of two matrices | The $L^2$ norm of the difference of the matrices | $\sqrt{\sum_{i,j=1}^k ([\Sigma_1]_{i,j} - [\Sigma_2]_{i,j})^2}$ |
| Centered Kernel Alignment (CKA) (Kornblith et al., 2019) | A normalized version of the Hilbert-Smith independence criterion (HSIC) (Gretton et al., 2005) between two sets of multivariate patterns | $\frac{HSIC_{U_1, U_2}}{HSIC_{U_1, U_1} HSIC_{U_2, U_2}}$<br>where $HSIC_{U_1, U_2} = vec(G_1)^T vec(G_2)$,<br>$G_1 = HU_1U_1^TH^T/p$, $H$ is the centering matrix, and $p$ is the number of columns of the corresponding stimulus-response matrix, $U_1$. |

It must be noted that by no means the list of metrics introduced in **Table 1** is exclusive. There are also other ways to quantify the similarity of two PSD matrices, but in this manuscript, we have considered the mostly popular metrics or those relevant to the Riemannian distance (e.g. the Frobenius norm of the difference).

### 2.2 An introduction to the Riemannian framework

This section introduces a Riemannian metric (Pennec et al., 2006; Pennec, 2006; Carmo, 1992; Bhatia, 2009) which has been used to measure distances between positive semi-definite (PSD) matrices with real elements.

We begin by introducing the general definition for the length of the curve that connects two PSD matrices. Then we explain how the relatively complicated equation describing this length could be simplified using the geometrical properties of the shortest curve connecting two PSD matrices calculated with the Riemannian metric.

Let *γ*(*t*) be a curve on the manifold of PSD matrices, connecting Σ_1_ to Σ_2_. When *t*, which is a variable, moves from 0 to 1, *γ* moves from *γ*(0) = Σ_1_ to *γ*(1) = Σ_2_ (**Figure 2**). The length of *γ* is defined as:

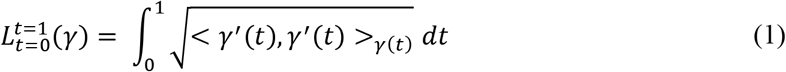

where *γ*′(*t*) is the derivative of *γ*(*t*) with respect to *t*, and <.,. >_*γ*(*t*)_ denotes the inner product defined on *γ*(*t*). The minimum-length curve on the manifold connecting any two arbitrary PSD matrices, here Σ_1_ and Σ_2_, is called the geodesic curve.

**Figure 2.**
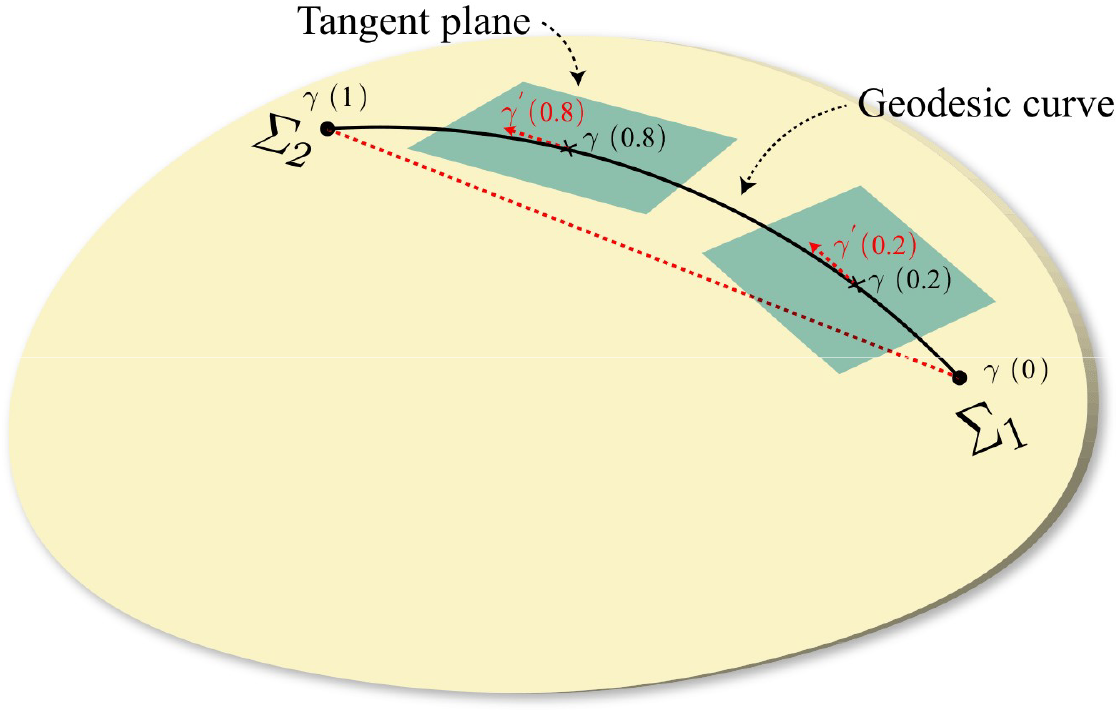
Riemannian manifold and geodesic curve. *This figure is a conceptual illustration for the geodesic curve, the tangent plane at a point on the manifold, and the difference between the straight line and geodesic curve. The geodesic curve showed in the figure connects Σ*_1_ = *γ*(0), *to Σ*_2_ = *γ*(1), *and its length can be calculated using Eq. 1 and Eq. 10. We also plot tangent planes at γ*(0.2) *and γ*(0.8)*. As we explained in the text and is also clear in the figure, γ*′*(t) lies in the tangent plane of γ*(*t*)*. This figure also shows that the straight line connecting Σ*_1_ *and Σ*_2_ *(shown with a dashed red line) and the geodesic curve could have different lengths. A critical point that we mentioned in the text is that the length of the geodesic curve is equal to the length of the line connecting Σ*_1_ *to the image of Σ*_2_ *on the tangent plane of Σ*_1_*. Notice that we have depicted the manifold as a 2D surface in* ℝ^3^*. However, the manifold of* 2 × 2 *PSD matrices cannot be shown in* ℝ^3^*, and hence this figure is supposed to just offer an intuition^4^*.

One way to quantify the distance between two PSD matrices, Σ_1_ and Σ_2_, would be to compute the length of the geodesic curve connecting them (**Figure 2**, black curve).

Two important properties of the geodesic curve help us derive a closed-form expression for its length. First, the norm of *γ*′(*t*), i.e. < 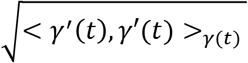, remains constant for all *t* ∈ [0,1]. Second, there is a unique geodesic curve between each pair of PSD matrices. The latter implies that the initial direction of the geodesic curve, *γ*′(0), could also be uniquely determined. Therefore, Eq. 1 can be reduced to:

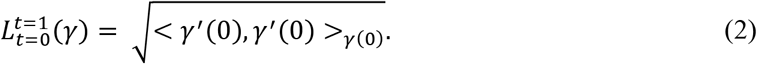

Hence, we only need the definition of the inner product and how to calculate *γ*′(0). In the following, we will explain these two important notions.

#### Riemannian metric

The Riemannian metric associates to each point (Σ) of the manifold an inner product <, >_Σ_ in the tangent plane of the point (for a detailed definition, see Carmo, 1992). The tangent plane at a point on the manifold is the hyperplane tangent to the manifold on that point (**Figure 2**, blue planes). It is clear that the derivative of *γ*(*t*), i.e., *γ*′(*t*), lies in the hyperplane tangent to the manifold at *γ*(*t*).

There can be an infinite number of choices for defining the inner product. Pennec et al. applied a constraint that leads to a unique definition. The constraint is that the metric and consequently the distances should remain invariant under an affine transformation. A metric with this characteristic is called an affine-invariant metric (Pennec et al., 2006):

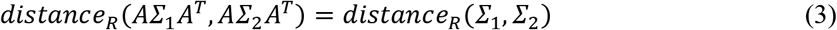

where *R* stands for Riemannian, *distance*_*R*_ is the length of the geodesic, and *A* is an arbitrary invertible matrix with the same size as Σ_1_ and Σ_2_. If we left-multiply a stimulus-response matrix by *A*, its 2^nd^-moment matrix will be in the form written in Eq. 3. In section 3.4, we discuss one potential application of this characteristic.

As we already explained, Eq. 1 can be simplified as the square root of an inner product, as in Eq. 2. Therefore, the affine-invariant property should be met at the definition of the inner product.

We start by defining the inner product at the point corresponding to the identity matrix on the manifold. A choice, inspired by the definition of the inner product in the Euclidean space, would be:

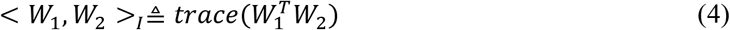

where *W*_1_ and *W*_2_ are two arbitrary matrices that lie in the tangent plane of the identity matrix, *I*. At the end of this section, it will be trivial that each matrix in the tangent plane is symmetric (Eqs. 7 and 8). The definition in Eq. 4, can be extended to the inner product at an arbitrary point Σ on the manifold using the affine-invariant property. Explicitly, Eq. 3 poses the following constraint on the inner product:

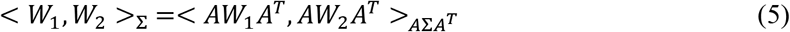

If we replace *A* with 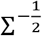, We will have:

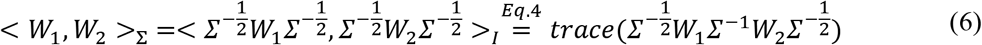

Having defined the inner product, we only need to calculate *γ*′(0) to compute the length of the geodesic (Eq. 1). For that, we need to define the notion of exponential and logarithmic transformations (these are also called exponential and logarithmic maps, (Pennec et al., 2006)).

We mentioned that in Eq. 1, *γ* is the geodesic curve that connects Σ_1_, *γ*(0), to Σ_2_, *γ*(1). As a special case, consider Σ_1_ = *I* and Σ_2_ = Σ.

*Exponential map.* It maps *tW* (*W* = *γ*′(0), and hence *W* ∈ tangent plane of *γ*(0)), to *γ*(*t*).

*Logarithmic map* (inverse of the exponential map). It maps *γ*(*t*) to *tW*.

As one can see, exponential and logarithmic maps are associations between the manifold and the tangent plane. Here, we introduce a mathematical expression for the exponential map and show this is the only possible choice. The proposed exponential map is:

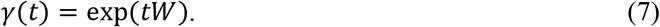

From linear algebra we have exp(*tW*) = *U diag*(exp(*td*_1_), …, exp(*td*_*n*_))*U*^*T*^, where *W* = *U diag*(*d*_1_, …, *d*_*n*_)*U*^*T*^ is the eigenvalue decomposition of *W* so that *UU*^*T*^ = *I* and *d*_*i*_ is the *i*^*th*^ eigenvalue of *W*. Since exp(*tW*) |_*t*=0_ = *I*, and exp′(*tW*)|_*t*=0_ = *W* and as we have already discussed, there is only one geodesic curve with *γ*(0) = *I* and *γ*′(0) = *W*. Hence, Eq. 7 is the only valid choice for the exponential map.

In order to derive *γ*′(0) we need to calculate log (Σ):

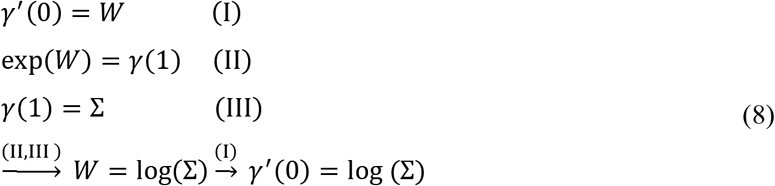

where (again from linear algebra we have that) log(Σ) = *Udiag*(log (*σ*_1_), …, log (*σ*_*n*_))*U*^*T*^.

Now the Riemannian distance between *I* and Σ can be derived. Also, the distance between any arbitrary Σ_1_ and Σ_2_ can be computed using the affine-invariant property, Eq. 3.

The closed-form equation for the Riemannian distance is as follow:

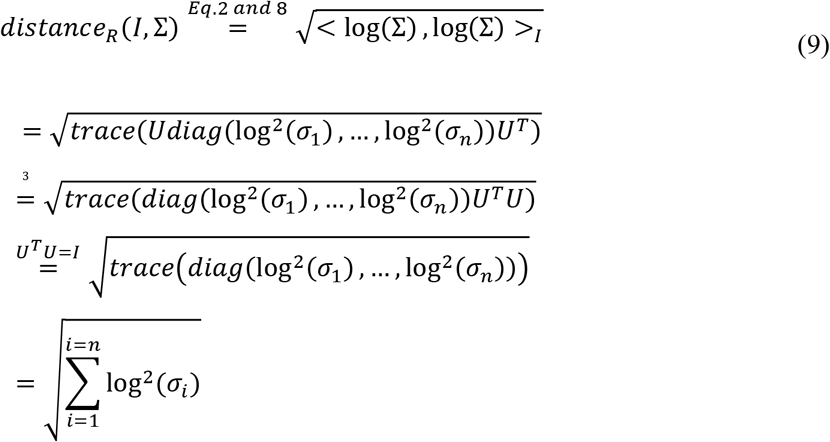

where *σ*_*i*_ is the *i*^*th*^ eigenvalue of Σ. From Eq. 3, we know that 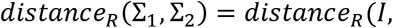 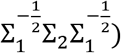. Hence, if we equate Σ in Eq. 9 with 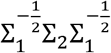, we will obtain the Riemannain distance between Σ_1_ and Σ_2_. Therefore, we have:

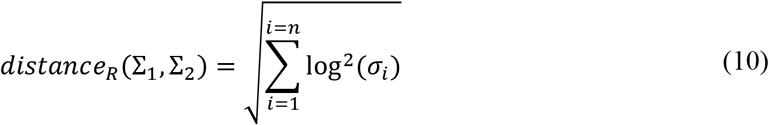

where *σ*_*i*_ is the *i*^*th*^ eigenvalue of 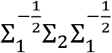, or equivalently the *i*^th^ eigenvalue of 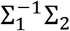.

From Eq. 8 we know that log (Σ) is the image of Σ on the tangent plane of *I*. The first line in 9 clearly shows that the length of the geodesic connecting *I* and Σ is equal to the norm of image of Σ in the tangent plane of *I*. Even more generally, one can show that the length of the geodesic connecting any arbitrary Σ_1_ and Σ_2_ is equal to the norm of image of Σ_2_ on the tangent plane of Σ_1_.

The above definition can be simplified for low-dimensional matrices (1 × 1 and 2 × 2). For example, in 1 × 1 matrices, we have:

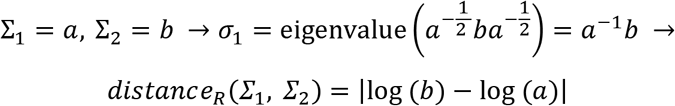

which is equal to the log-distance of *a* and *b*.

#### Riemannian Mean

Similar to the Euclidean mean, the mean of a number of PSD matrices could be defined considering their underlying manifold. This is often referred to as the Riemannian mean (Pennec, 2006). Although we do not use this notion in this manuscript, we include the definition here as there are potential applications for it in future studies.

Given *n* points ∈ *sym*^+^, the Riemannian sample mean (Σ_*m*_) is defined by:

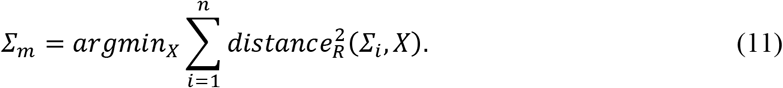

We can see that for the 1-D case, this definition yields the geometric mean.

### 2.3 Providing meaningful zero point for distance-based metrics using condition-label permutation

When using correlation-based metrics for comparing two representational structures, non-duplicative elements - lower triangular elements in case of RSM or lower triangular plus diagonal elements in case of 2^nd^-moments - are vectorized, and the correlation between two vectors is calculated. It is expected that the correlation of two matrices that are not related would be zero. Therefore, a zero value for the correlation metric implies that the representations are not related. Hence, this meaningful zero point allows us to benefit from (non-)parametric statistical tests that can validate different types of hypotheses investigated in RSA (this point will be discussed in the Results section). In order to provide such a meaningful zero point for distance-based metrics, we employ a type of non-parametric test called the permutation test (Nichols and Holmes, 2002).

In the permutation test, the distribution of the test statistic under the null hypothesis is obtained by calculating the values under all (or a large number of) possible equiprobable rearrangements. **Figure 3** shows an example of rearranging two conditions of an RSM. Here we leave one RSM unchanged and consistently permute the rows and columns of the other RSM for a large number of times and compare them in each iteration. The comparisons obtained under different permutations constitute the null distribution. If two RSMs have similar structures, we expect a few numbers of the null estimates to be more extreme than the summary statistic obtained under neutral permutation. Note that being more extreme translates differently for distance-based and correlation-based metrics. For distance-based metrics, it refers to smaller values, and for correlation-based metrics, it refers to larger values. Therefore, we can obtain a p-value for testing the similarity of two RSMs for each metric. The p-value is the proportion of values in the null distribution that are more extreme than the summary statistic. Smaller p-values indicate RSMs that are more similar according to a metric.

**Figure 3.**
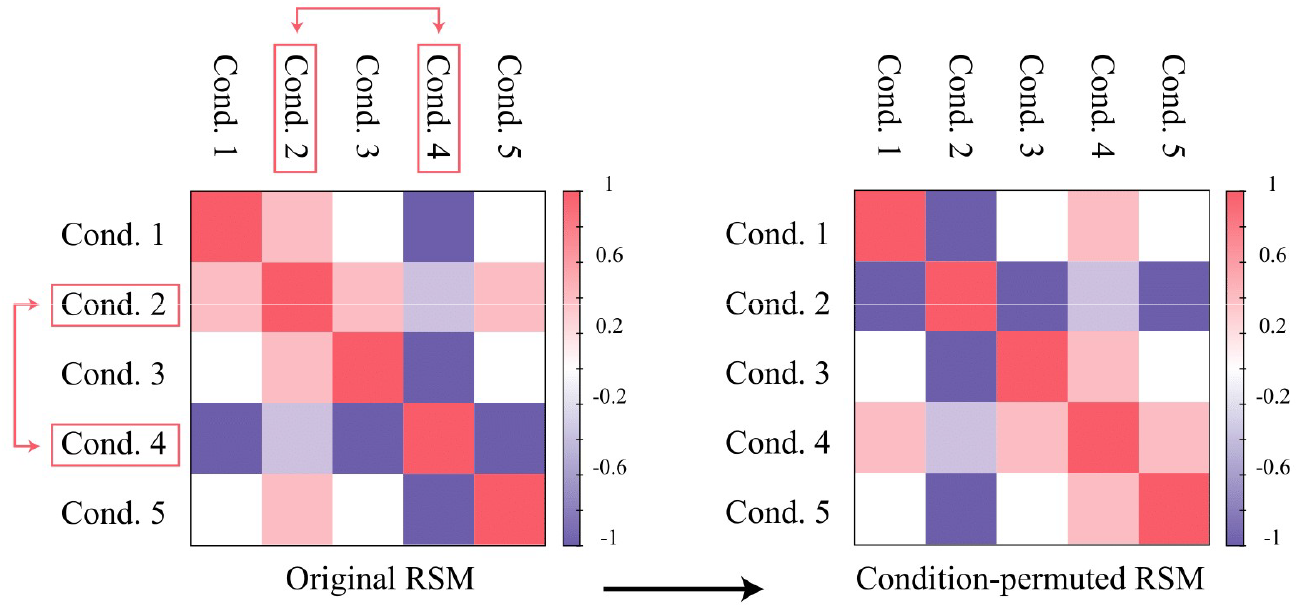
Permutation of condition-labels. *This figure illustrates a simple example of condition permutation in a* 5 × 5 *RSM, where the second and the fourth conditions corresponding to the second and fourth rows and columns are replaced with each other. For a k* × *k RSM, there can be k*! *condition-permuted matrices, all of which are obtained by similarly rearranging rows and columns. However, it is not necessary to exhaustively try all different rearrangements.*

Now we can have the bias-correlated distance (correlation) of two RSMs by subtracting the summary statistic from the mean of the null distribution (mean of the null distribution from the summary statistic).

The bias-corrected metrics are expected to have a zero mean under the null hypothesis. Hence, a meaningful zero point that allows for making inference can be achieved. Note that assuming a theoretical mean on the null distribution (e.g., zero correlation under the null) obviates the need to estimate the empirical null. However, for some metrics, such as the Riemannian distance, which are distance-based and have non-negative values, the only way to make inference would be to eliminate the bias for each estimated value.

By our definition, all bias-corrected metrics, correlation-based or distance-based, estimate similarities, meaning that a significant positive deviation from zero implies similarity between RSMs. Henceforth, we use bias-corrected metrics to estimate similarities (or relatedness) between different representational matrices, to which we refer as *similarity*_*bc*_, where bc stands for bias-corrected.

## 3 Results

### 3.1 The Riemannian distance captures representational relationships more accurately when there are relatively small number of response channels (e.g. voxels)

Each measurement modality provides an estimate of the brain activity for a number of response channels. These response channels are voxels for fMRI, electrodes for M/EEG, and cell recordings for electrophysiological data. Generally, the larger the number of response channels per a brain region, the more accurate the estimates of its response properties. Nonetheless, there is often a sampling bias in measuring brain responses (Panzeri and Treves, 1996). In this part, we focus on the effect of that bias when dealing with 2^nd^-moment matrices and RSMs. Specifically, we explore whether different metrics have different levels of susceptibility to the sampling bias.

Similar to encoding analysis, we treat activity profiles (vector of activities for a single voxel) for a number of conditions as points in the space spanned by experimental conditions (Diedrichsen and Kriegeskorte, 2017). Therefore, the 2^nd^-moment matrix can be obtained from those point clouds (one point per voxel). Having that in mind, with a larger number of response channels, we would have richer sampling and a more accurate estimate of the 2^nd^-moment matrices and thus the RSMs. However, in almost all cases, there is an inevitable sampling bias. Therefore, if a similarity measure can afford capturing the relationship between RSMs with few neurons/voxels, that will be count as an advantage for it. Additionally, in many cases the number of neurons/voxels are dependent on the choice of the experimenter and it would be best if these decisions have little influence on the results^5^.

**Figure 4** shows two sets of response patterns (different colors) for two experimental conditions (each axis corresponds to one condition). Each point in each panel corresponds to one voxel. While the two distributions are clearly different in **Figure 4**A, reducing the number of response channels can make it difficult to distinguish different structures. With a small number of samples (**Figure 4**B) one cannot discern between the two representational structures. However, richer sampling (**Figure 4**A) correctly concludes that the two are distinct.

**Figure 4.**
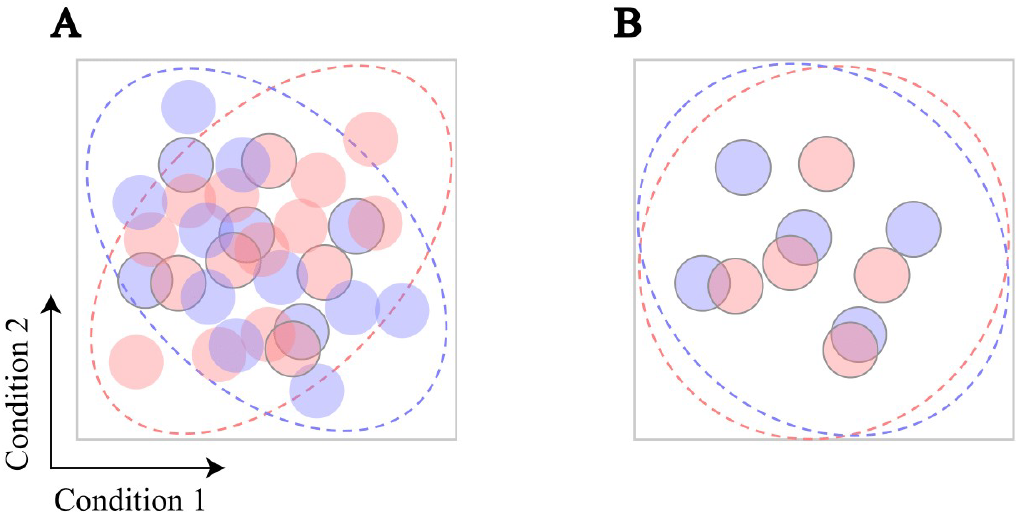
Richer sampling of the condition space allows for adjudicating between different representational models. Consider the responses of two brain regions to two different stimuli. We can show the responses as points in the condition space (each color corresponds to one region). **A.** With a large sample size, it is evident that the two regions have distinct distributions, hence different 2^nd^-moment matrices. **B.** However, with only a few samples, the distributions of activity profiles may not be distinct.

We use simulations with known underlying representational structures (i.e., RSMs or 2^nd^-moments depending on the analysis) to compare different metrics in terms of their dependencies on the number of response channels. Particularly, we evaluate the performances of these metrics using two different tests, namely, the *relatedness* and *comparison* tests. The former examines whether representational structures are statistically *related* or not, and the latter compares the similarity of different representational structures to a reference. In the following, without loss of generality, we use RSM to refer to a representational structure. Additionally, since what we describe here is general and not specific to fMRI data, we will use voxel and response channels interchangeably.

In each simulated subject, for a given number of conditions we generate two sets of response patterns, each containing 1000 samples from zero-mean multivariate Gaussian distributions with known distinct covariance matrices, *R*_*a*_ and *R*_*b*_ (**Figure 5**). We then derive various RSMs from each set of patterns by randomly sampling from the voxels (for example, 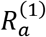, …, 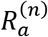 in **Figure 5**). Intuitively, we expect that for a large number of samples (i.e., voxels), the new RSMs would be similar to their corresponding RSMs (**Figure 5**, dashed black arrow). Therefore, we expect the variants of the original RSMs to be significantly *related* to the original ones. Additionally, since the two underlying structures are distinct, we expect the new variants from each to be closer to themselves than to the two variants of the other structure (for example, in **Figure 5**, 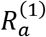 should be more similar to 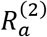 than to 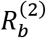.

**Figure 5.**
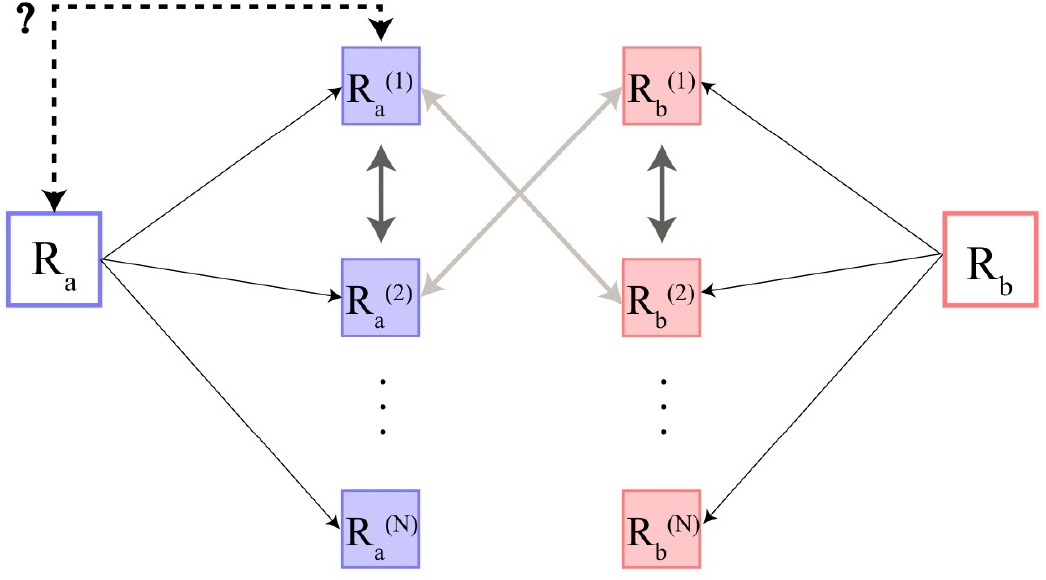
Simulation setup for evaluating dependency of metrics on the number of response channels (e.g., voxels). Here R_a_ and R_b_ are two distinct RSMs that serve as ground truths for simulations. From each, we obtain response patterns with a large number of voxels. 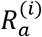 and 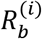 are RSMs derived by sampling the voxels from each set of patterns. We prefer a metric for which, 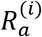 is closer to 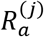 than to 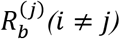. For example, blue-to-blue arrows should show stronger similarity than red-to-blue arrows. Furthermore, we prefer a metric for which 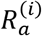 is “related” to the ground truth (R_a_). For example, the relationship depicted by the dashed arrow is preferred to be statistically significant.

#### Testing for relatedness

For each 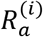 we perform a condition-label permutation test (as explained in section 2.3) to obtain a p-value for testing the *relatedness* of *R*_*a*_ and 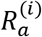. Additionally, we repeat this for a large number of iterations (subjects) and then count the proportion of significant tests after controlling the expected FDR (Benjamini and Hochberg, 1995) at 0.01. This score would be a measure of *significance mass*, conceptually similar to what is explained in (Allefeld et al., 2016) and we use it to compare different similarity measures. Intuitively, we prefer a measure that can detect the relatedness despite a having small number of samples (voxel counts).

**Figure 6** shows the proportion of significant tests as a function of the number of voxels for different metrics. As it can be seen, the Riemannian distance shows strong superiority in comparison to others. For 2^nd^-moment matrices, linear CKA also gives intermediate values that are better than the rest of the metrics. Note that linear CKA can only be used to quantify the relationship between 2^nd^-moment matrices and thus it would not be applicable to RSM comparisons. For obtaining the results of **Figure 6**, we used the Visual object recognition dataset (Hanson et al., 2004; Haxby et al., 2001; O’Toole et al., 2005) to define *R*_*a*_; however, the results are robust to the choice of the ground truth. In fact, the standard error in **Figure 6** is calculated across different choices of *R*_*a*_. We used the data from 10 visual ROIs (V1-V5, left and right) per subject for *R*_*a*_.

**Figure 6.**
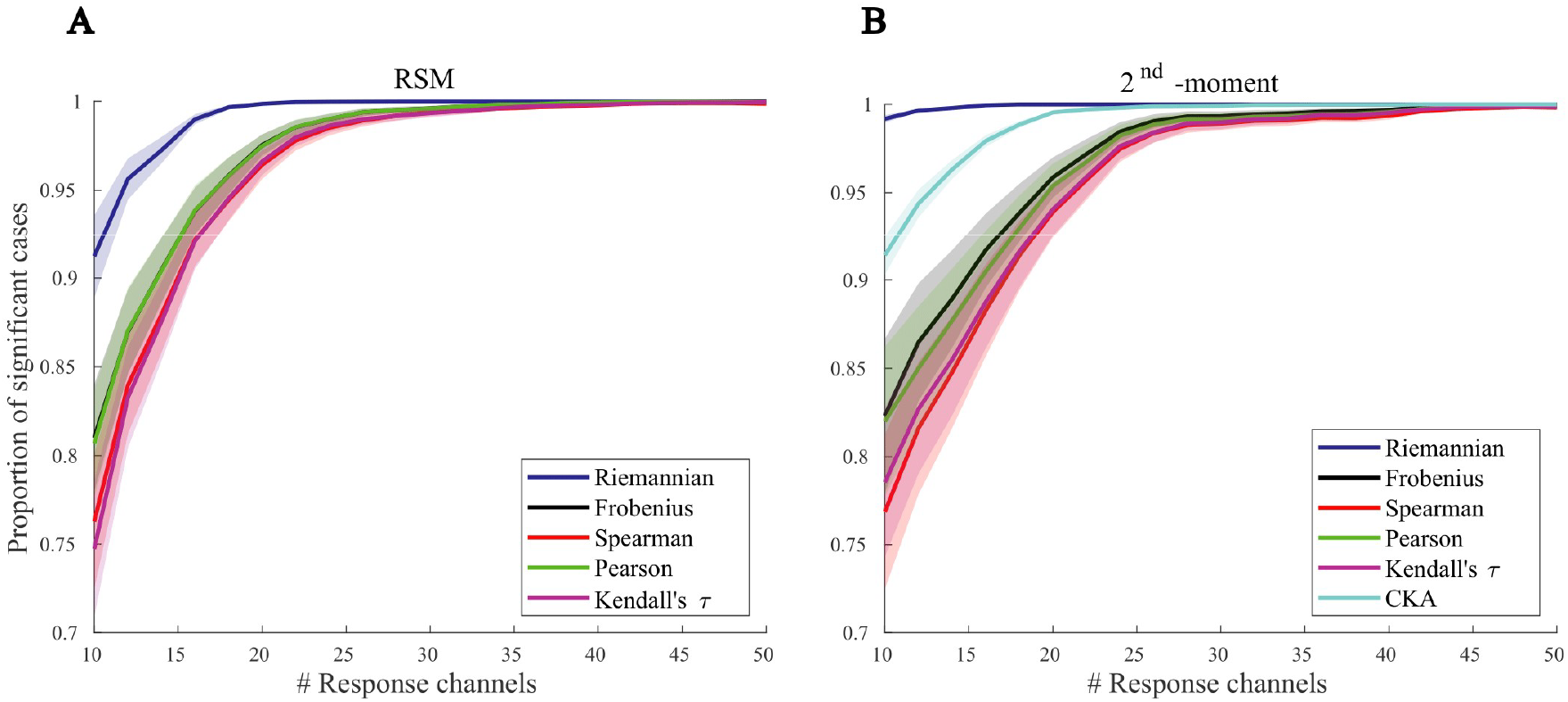
Evaluating the dependency of metrics to the number of response channels when testing for “relatedness”. For each number of response channels, points of the curves are the proportion of significant cases in which the relatedness of R_a_ and each of the 20 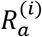 s is tested. We repeated the simulation for 10 (V1-V5, left and right) R_a_s to obtain the standard error. **A.** Results for RSMs. The Riemannian distance shows the best performance among metrics. **B.** Results for 2^nd^-moments. The Riemannian distance has the best performance and the CKA show the second-best performance.

#### Comparison test

The RSMs from two groups *a* and *b* would be discriminable (Nili et al., 2020) if the following inequalities hold: (**Figure 5**).

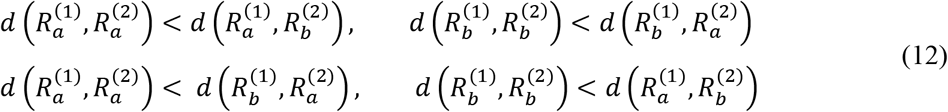

Here we have used the distance notion; however, one can use the correlation notion with reversed inequalities. Further, same inequalities should hold for 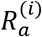, 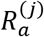, 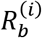, and 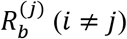. For distance-based metrics we can define a score as:

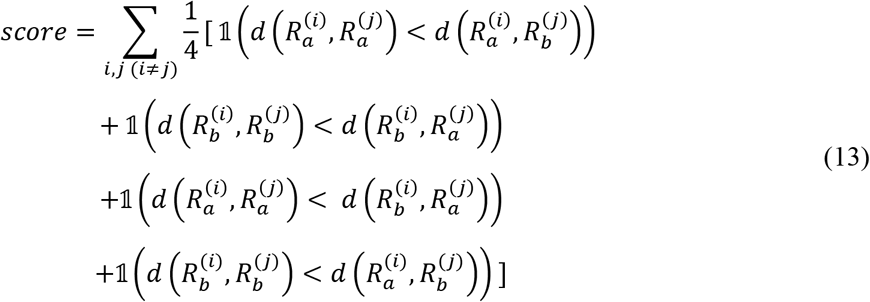

The score will be one if all the inequalities hold. For each voxel count, we calculate the score multiple times by repeating the simulation for different choices of *R*_*a*_ and *R*_*b*_ and take the average. A metric that has a higher average score is superior as it distinguishes variants of two primarily distinct RSMs with more sensitivity.

**Figure 7** shows the scores for different metrics. In this simulation, we use Visual object recognition dataset to obtain *R*_*a*_ and *R*_*b*_ (**Figure 5**). Scores are calculated for a range of voxel counts. Results are presented separately based on the type of the representational structure (RSM or 2^nd^-moment). Whether RSMs are used or 2^nd^-moment matrices, in almost all cases the Riemannian distance has an outstanding superiority. When RSMs are examined, all metrics except the Riemannian distance, which gives the best score, show similar performances. When 2^nd^-moment matrices are examined, again the Riemannian distance has the highest score and linear CKA has the second highest score. As a control measure, the Frobenius norm (another measure of the distance between two matrices) behaves poorly in comparison with other metrics (**Figure 7**B). So, the superiority of the Riemannian metric cannot be attributed to its distance-based nature.

**Figure 7.**
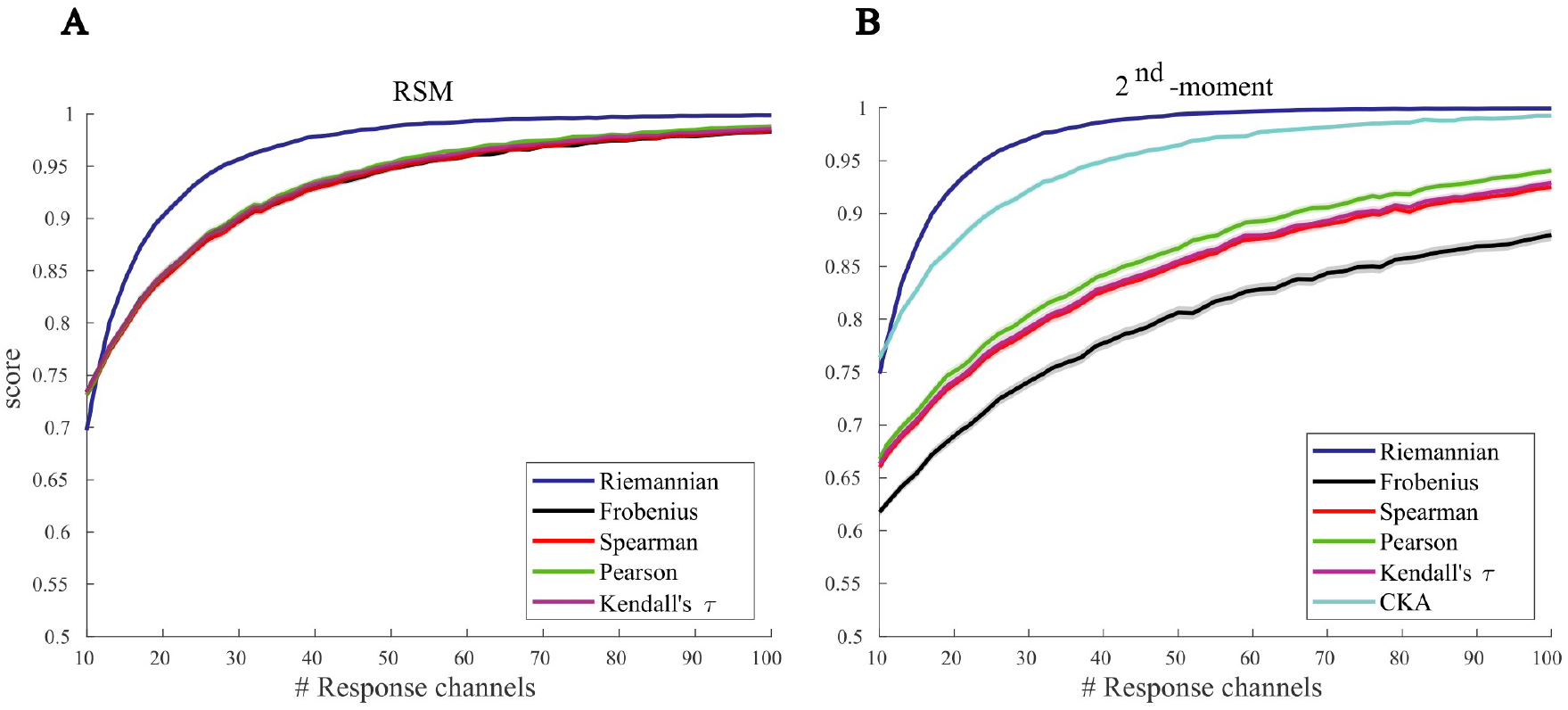
Evaluating the dependency of metrics to the number of response channels in the comparison test. For each number of voxels, curves show the scores (Eq. 13) for different metrics. The simulation has been repeated for all pairs of visual regions of the brain of an exemplar subject as seeds, and the average score is plotted. Error bars show the standard error. **A.** Results for RSMs. Apart from CKA, the Riemannian distance shows a clear superiority, and other measures behave similarly. **B.** Results for 2^nd^-moment matrices. Again, the Riemannian distance has the highest average scores.

### 3.2 Comparing representational structures via Riemannian distances yields higher intra-subject consistency (within-subject reliability)

In this section, we compare different metrics in terms of their reliabilities. Intuitively, a metric that results in larger test-retest reliability would be preferred (Walther et al., 2016). In order to quantify the reliability of different metrics, we compare representational structures (RSMs and 2^nd^-moment matrices) for a subject in different functional runs and make across-run comparisons in a range of ROIs. The across-run comparisons are summarized by a measure that we refer to as the *consistency score*. The consistency score determines the degree to which independent instances of representational structures, derived from different functional runs, are significantly related.

For a subject, we define the consistency of a region’s representations across runs as:

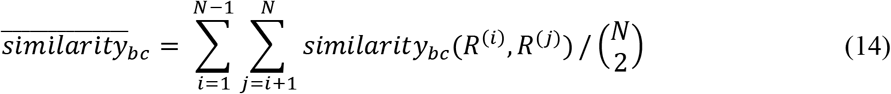

where bc stands for bias-corrected (section 2.3), *N* is the number of runs, and *R*^(*i*)^ is the representational matrix of the *i*^*th*^ run. In other words, 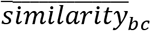 is the sample mean of bias-corrected values (distances or correlations), computed between RSMs or 2^nd^ moment matrices of all pairs of runs in an ROI. The consistency score for this region is the t-statistics of 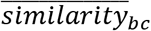s(tested against zero) calculated across all subjects. (**Figure 8**).

We use two datasets for this analysis: Visual object recognition dataset, and Generic object decoding dataset, and show results for different metrics (including the Riemannian distance). When RSMs are used as representational structures (left column in **Figure 9**), the Riemannian distance shows a higher degree of within-subject reliability compared to other metrics. This can be deduced by the observation that the consistency scores (pooled across ROIs) of the Riemannian distance have a statistically significant larger median than the consistency scores of almost all other metrics. Each solid black line indicates the significance of the paired test (Wilcoxon signed-rank test) of consistency scores, obtained from multiple ROIs, with the Riemannian distance and another metric. These results are also replicated for 2^nd^-moments (right column in **Figure 9**).

**Figure 8.**
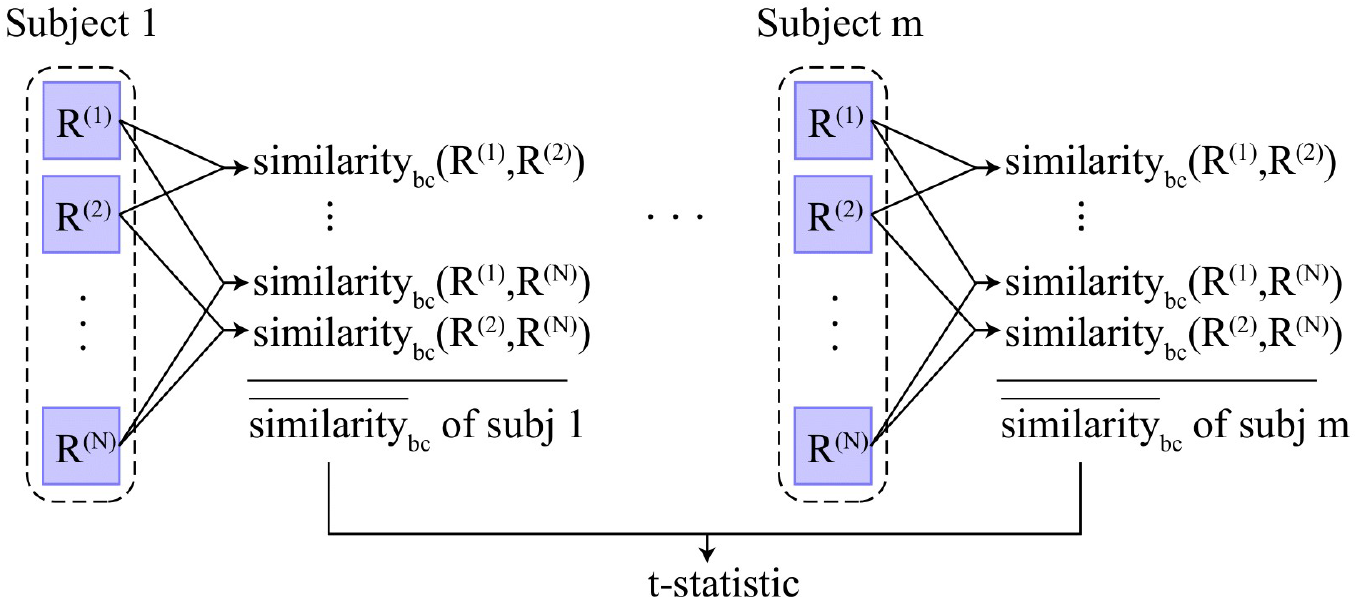
Calculation of the consistency score. First, for each subject, the average of bias-corrected values (distances or correlations) between RSMs of all pairs of runs (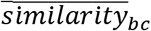 Eq. 14) is calculated. Then, the consistency score is calculated as the across-subject t-statistic of those values. This score can then be pooled across ROIs and tested against zero or be compared for different metrics. We say that one metric is more consistent than another if it results in larger consistency scores.

**Figure 9.**
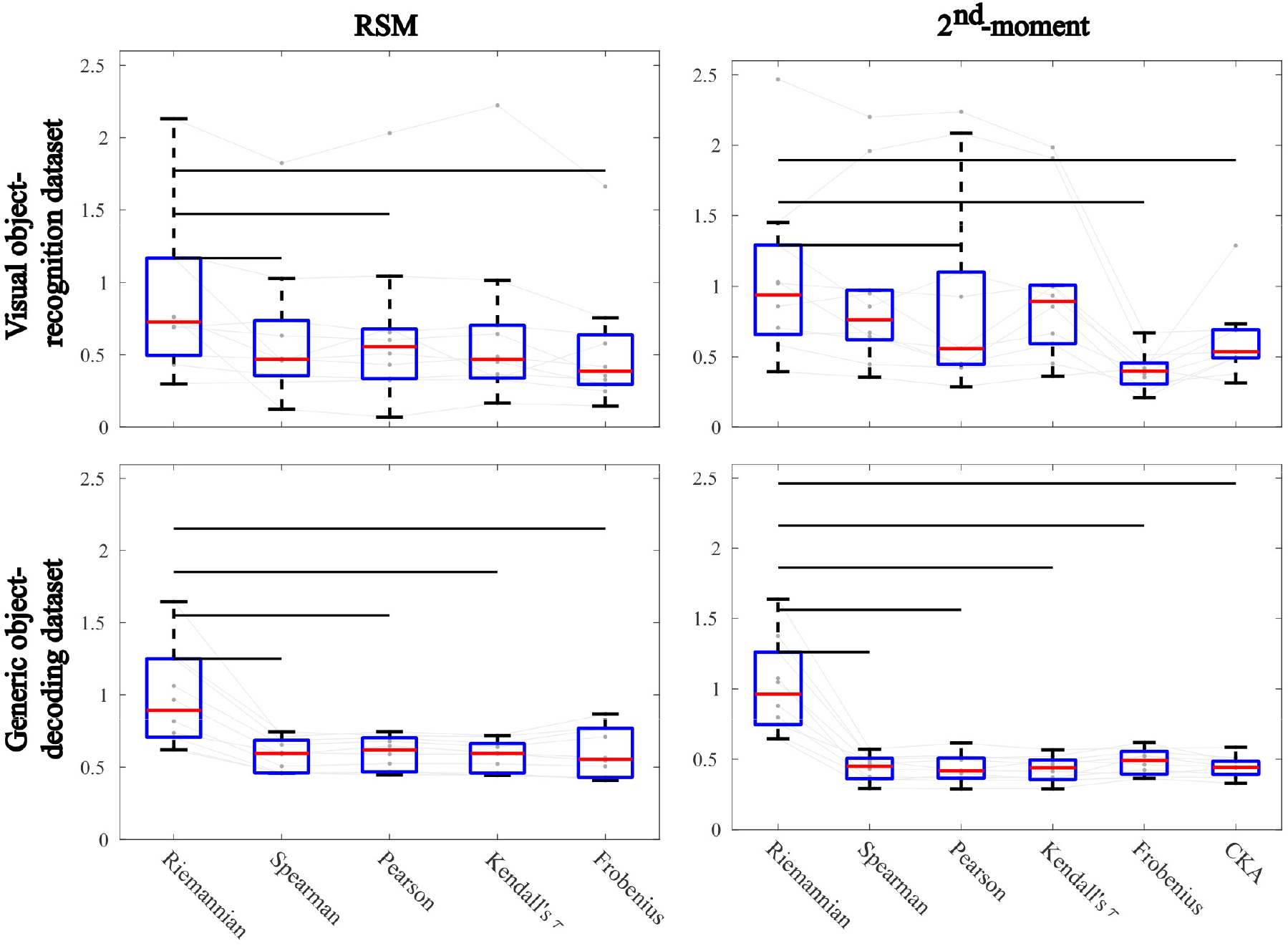
Comparing within-subject reliability (consistency) of different metrics. Results for the Visual object recognition and Generic object decoding datasets are shown in the first and second rows, respectively. Furthermore, different columns correspond to results from RSMs (left) or 2^nd^-moment matrices (right). For each panel of the figure, if there is a horizontal black line between the Riemannian distance and each of the other metrics, it means that the consistency scores (pooled across ROIs) attributed to the Riemannian distance has a statistically larger median (p<0.05, Wilcoxon signed-rank test) than the consistency score of another metric. Each slim gray line appearing at the background of the figure shows the consistency score of one ROI across different similarity metrics.

### 3.3 Comparing representational structures with the Riemannian distance yields higher discriminability for representations from distinct brain regions

Here we explore the extent to which distinct representations are discriminable according to various measures of representational similarity. To this end, we define a summary statistic called the *discriminability score*.

The discriminability score determines if independent instances of RSMs (or 2^nd^ moment matrices) from two distinct brain regions are discriminable.

We define the discriminability score for regions *a* and *b* in a subject as:

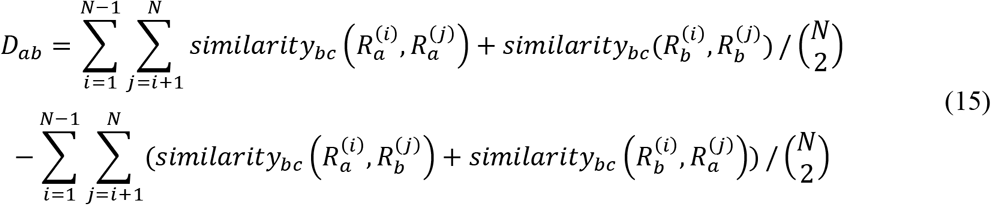

where *N* is the number of runs, and 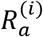 and 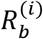 are the representational matrices of the *i*^*th*^ run of the regions *a* and *b*, respectively. The details for computing the discriminability score is depicted in **Figure 10**. In the figure, *D*_*ab*_ is the difference between the average of darker arrows (bias-corrected distances or correlations of the samples of RSMs or 2^nd^-moment matrices from the same region) and lighter arrows (bias-corrected distances or correlations of the samples from distinct regions). If *D*_*ab*_ is large, the darker arrows in **Figure 10** show substantially larger similarities than the lighter arrows. The discriminability score for regions *a* and *b* is the t-statistic (tested against zero) of *D*_*ab*_s calculated across all subjects.

A metric has a significantly larger discriminability score than another one if its discriminability scores, pooled across all pairs of studied regions, are significantly larger than the scores of another metric. We test this hypothesis using the one-sided Wilcoxon signed-rank test.

**Figure 10.**
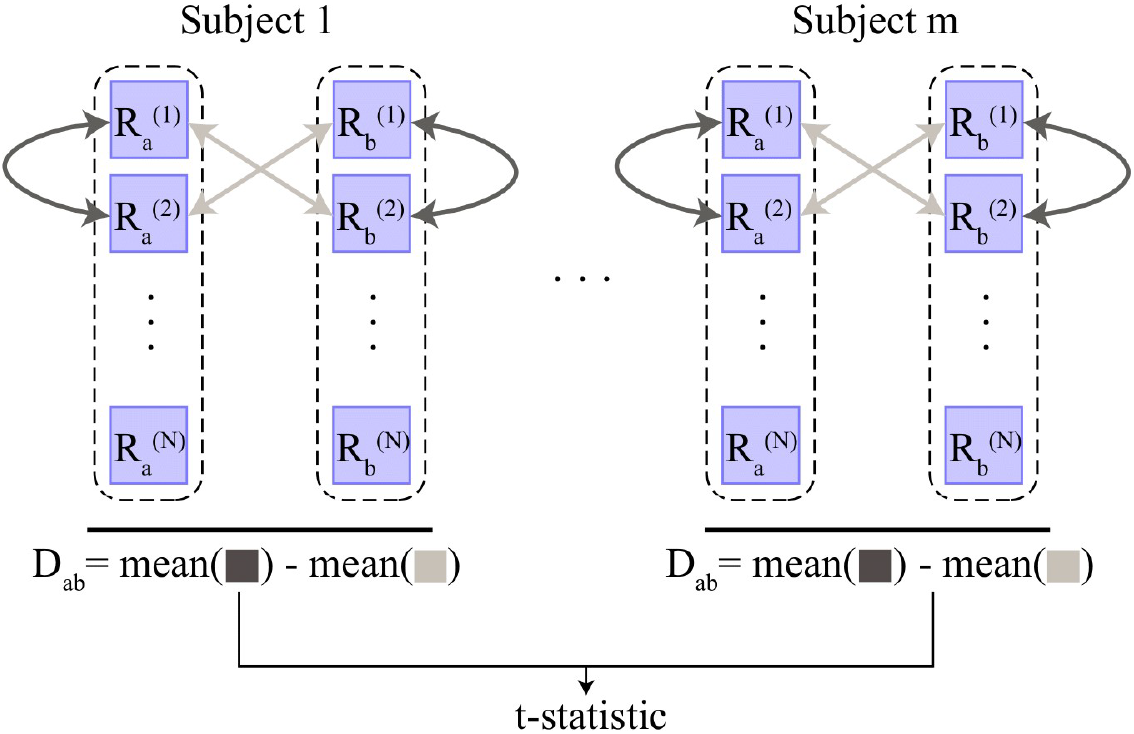
Calculation of the discriminability score. For regions a and b of each subject, the difference between the average of the bias-corrected values between RSMs from the same region (darker arrows) and the average of the bias-corrected values between RSMs from distinct regions, D_ab_ (Eq. 15), is calculated. The discriminability score for regions a and b is the t-statistic (tested against zero) of D_ab_s across subjects. This score can then be pooled across all pairs of regions and compared for different metrics. We say that one metric shows a higher degree of discriminability if it results in larger discriminability scores on average. This can be tested by a one-sided Wilcoxon signed-rank test.

Similar to the previous section, we used Visual object recognition dataset, and the Generic object decoding dataset. In either case where RSMs (first column in **Figure 11**) or 2^nd^-moment matrices (second column in **Figure 11**) are used as the representational structure, the Riemannian distance performs best in discriminating representations from non-overlapping regions. This can be deduced by the observation that the discriminability scores (pooled across pairs of ROIs) of the Riemannian distance have a statistically significant larger median than the discriminability scores of other metrics. Solid black lines indicate the significance of the paired tests.

**Figure 11.**
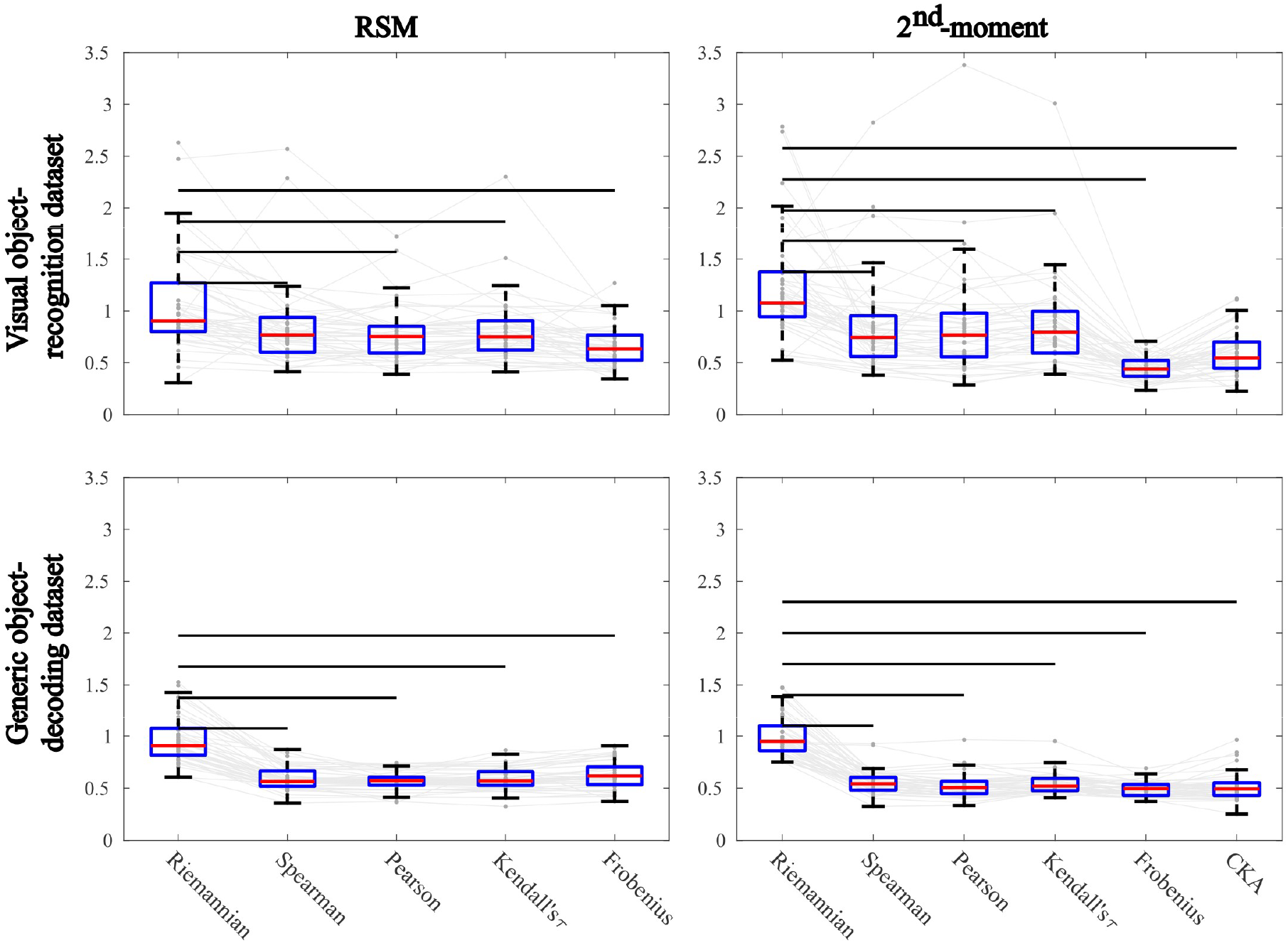
Comparing discriminability scores across metrics. The Visual object recognition dataset (first row) and the Generic object decoding dataset (second row) are used in our analysis. Column determines whether RSMs or 2^nd^-moment matrices are used as representational structures. For each panel of the figure, if there is a horizontal black solid line between the Riemannian distance and each of the other metrics, it means that the discriminability scores (pooled across pairs of regions) attributed to the Riemannian distance has a statistically significant larger median (p<0.05, one-sided Wilcoxon signed-rank test) than the discriminability scores of another metric. Each slim gray line appearing at the background of the figure, shows the discriminability score of a pair of ROIs across different metrics.

**Figure 12.**
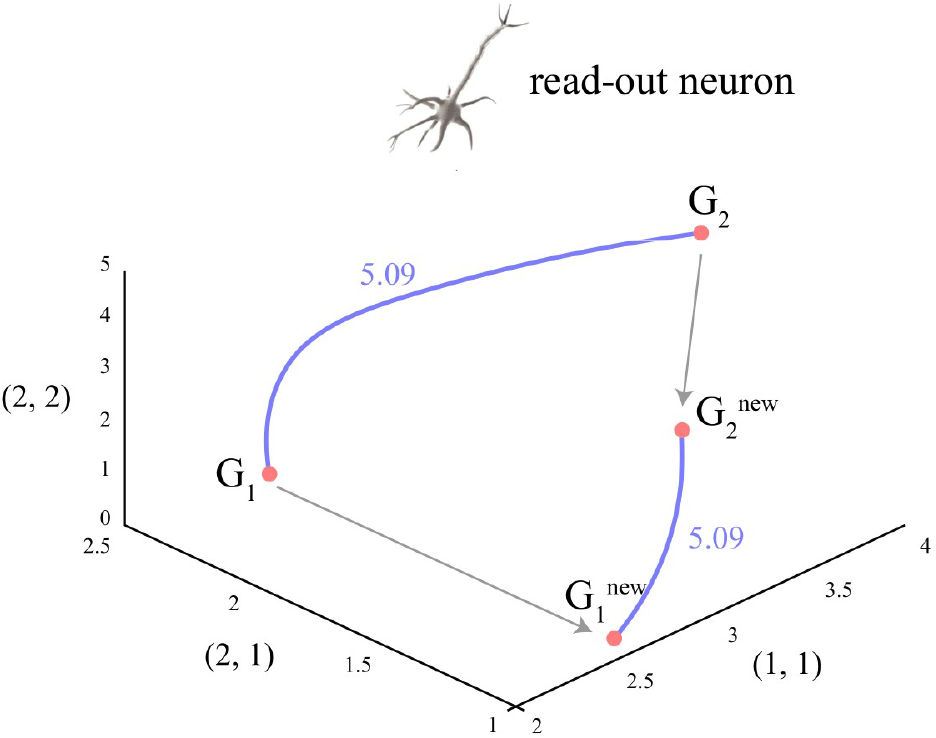
Identical full-rank linear operations on activity patterns of two regions does not affect their representational similarity when the Riemannian distances are used. *The figure depicts two* 2 × 2 *2^nd^-moment matrices, G_1_ and G_2_, and their transformed versions, G_1_^new^ and G_2_^new^. The length of the geodesics (solid blue curves) that connect the 2^nd^-moment matrices before and after linear transformations are identical (5.09 in this case) whereas their Euclidean distance changes.*

### 3.4 Motivations for using the Riemannian distance when testing for shared information: identical loss less linear transformations of activity patterns from two regions preserves their relationship

Given the benefits we saw in using the Riemannian distance for comparing representations, we next explore the advantages of using it for testing shared information or representational connectivity (Basti et al., 2020). In this section, we explain a compelling property of the metric that makes it particularly useful for testing shared information.

Multi-dimensional connectivity methods quantify the shared information between two sets of response patterns. Multi-dimensional connectivity can be detected and exploited via a read-out neuron. A read-out neuron that has access to both regional patterns can reveal the shared information by applying a set of weights to the activity patterns. For example, Informational Connectivity (Coutanche and Thompson-Schil, 2013), the first method of this class, establishes functional connectivity if classification accuracies correlate across time in two separate regions. This would imply shared information about a particular binary distinction. Notably, this information can be detected and exploited by the read-out neuron.

Importantly, it is plausible that the read-out neuron mixes the activity patterns in the two regions (or receives the mixed patterns) in the same way. Therefore, we would favor a metric that does not change with identical lossless mixing of the response patterns. Mathematically, if we denote the stimulus-response matrices of the two regions as *U*_1_ and *U*_2_, where *U*_1_ is a *k* × *p*_1_ matrix (stimulus × channels) for one region and *U*_2_ is a *k* × *p*_2_ matrix of responses to the same stimuli in another region, we want to have:

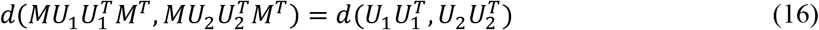

conditioned on *M* being a full-rank (*k* × *k*) matrix.

Interestingly, among all different metrics, the Riemannian distance is the only one that satisfies this property^6^.

To illustrate this with an example, consider two representational spaces (e.g. two searchlight ROIs), with 2^nd^-moment matrices G_1_ and G_2_. These two matrices may correspond to two points in the space of the PSD matrices. Now, applying the same mixing matrix, *M*, to the activity patterns of the two regions, results in two new 2^nd^-moment matrices: G_1_^new^ and G_2_^new^. These two matrices might have different distances in the Euclidean space. In other words, the length of the straight lines that connect two given matrices changes due to the mixing operation. However, when considering the length of the geodesic curve that connects two matrices on the Riemannian manifold, the distance between them (and hence their relationship summarized by the Riemannian distance) is unchanged after the transformation.

This is a desirable property as identical full-rank linear transformations are applied to the patterns in the two regions and intuitively this should not affect their shared information.

### 3.5 Advantages of using Riemannian distances when testing for multi-dimensional connectivity: detecting non-linear regional interactions with high sensitivity

Representational similarity of representations from different brain regions, implies shared information, functional connectivity, and representational connectivity (Basti et al., 2020; Kriegeskorte et al., 2008). In this section, we explore the advantages of exploiting the Riemannian distance for quantifying representational connectivity. To this end, we compare the Riemannian distance with measures introduced earlier for representational connectivity: RCA (Basti et al., 2020; Kriegeskorte et al., 2008), distance correlation (dCor; Székely & Rizzo, 2014), and CKA (Kornblith et al., 2019). Since other measures discussed in this section have reserved abbreviations, we have adopted dRiem for the Riemannian distance in the following and the next section.

It has been argued that 2^nd^-moment matrices fully capture the representational content of brain activity patterns (Diedrichsen and Kriegeskorte, 2017). For that reason, when comparing representations in two brain regions we compute the Riemannian distance between their 2^nd^-moment matrices^7^. However, that comes with a cost as it requires the 2^nd^-moment matrices to be positive definite. This means that the number of conditions needs to be less than the number of voxels, which we observe in our simulations.

Similar to (Basti et al., 2020), we consider simulated examples in which two regions are functionally connected. In these cases, it is expected that measures capture the functional connectivity.

Let *X* and *Y* be the response patterns of two regions (condition × voxel matrices). We have:

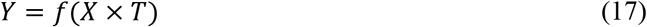

In this case *T* is a matrix that maps the voxels in one region into another. Additionally, *f* can be an arbitrary function. We take *X* from a multivariate normal distribution and use a random matrix for the mapping matrix, *T*. Both *X* and *Y* are contaminated by multivariate Gaussian noise.

#### Multivariate linear relationship

As a proof of concept, we first tested this simple linear connectivity scenario. In this case, *f* is the identity operator, thus *Y* = *X* × *T* with some additive noise. We simulated data for a number of *subjects* (N=50) and two regions. Furthermore, we explored different combinations for the number of voxels and the number of conditions, observing the constraint that the number of voxels be larger than the number of conditions. For each set of parameters, we estimated different connectivity measures and their null values for a number of simulated subjects (section 2.3). We then obtained the bias-corrected measures and their corresponding across-subject z-scores.

**Figure 13** shows the z-maps of different metrics for various simulation settings. Clearly, all measures can capture the simulated *functional connectivity* robustly and the z-scores are all highly significant.

**Figure 13.**
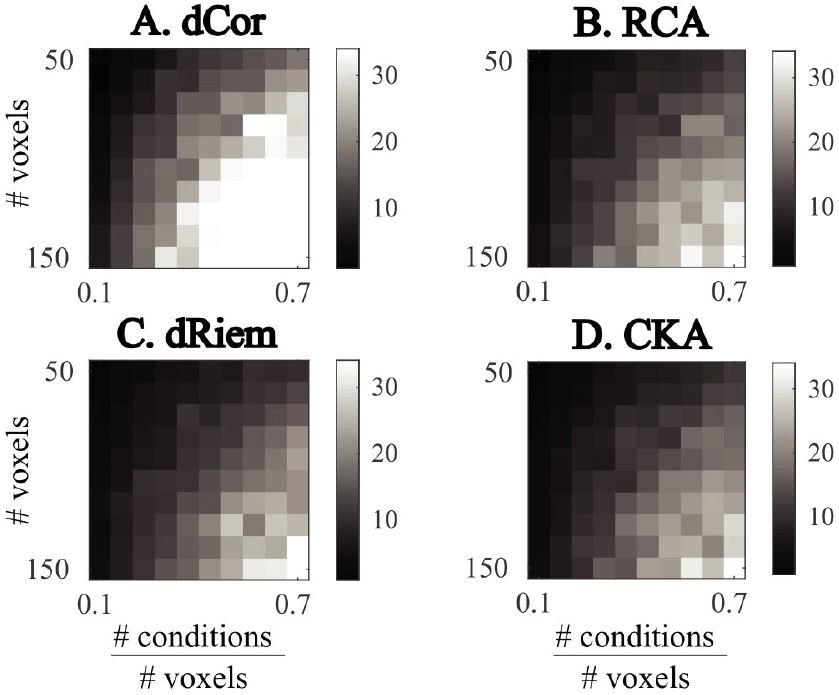
Comparing different measures of representational connectivity for simulated data with multivariate linear connections. *We simulated 50 subjects. For each subject a stimulus-response matrix in one region, X, was obtained by sampling from a multivariate normal distribution. The stimulus-response matrix in the other region, Y was X* × *T, with T being a constant voxel-space transformation. Each element of T was chosen randomly from a normal distribution. X and Y were then contaminated by i.i.d samples of additive white Gaussian noise. Panels A-D represent the z-scores of bias-corrected connectivity measures calculated across subjects. As one can see, in almost all voxel-condition counts, all metrics could detect the linear association between regions (z*>*1.645)*.

Similar to the previous sections, we could have also computed Riemannian distances between RSMs instead of 2^nd^-moment matrices. The reason we used 2^nd^-moment matrices is three-fold. First, RSMs could be directly derived from 2^nd^-moment matrices and thus the similarity of 2^nd^-moment matrices implies similarity of RSMs^8^. Second, the 2^nd^-moment matrices contain information about the overall activations as well. The diagonal elements define the *L*_2_ norm of the activation for different conditions. Therefore, assessing representational connectivity from the similarity of 2^nd^-moment matrices is a more comprehensive test that also incorporates univariate connectivity. Third, our simulations confirm that sensitivity is higher when 2^nd^-moment matrices are tested (**Figure 14**).

**Figure 14.**
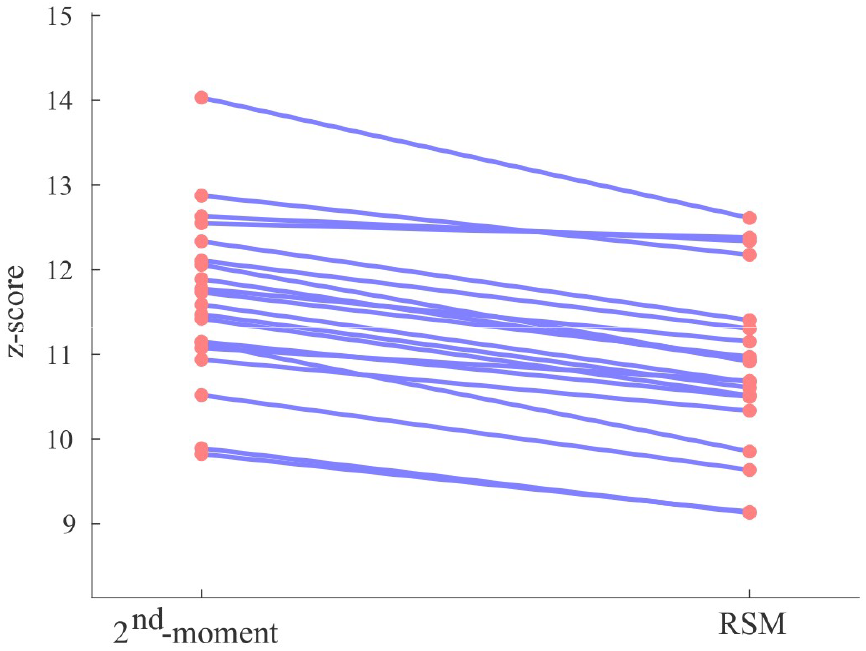
The Riemannian distance between 2^nd^-moment matrices of two linearly connected regions gives larger effect sizes (z-scores) than the Riemannian distance between their RSMs. For 20 subjects, we simulated two linearly connected regions, with a condition number of 30 and a voxel count of 40, and corrupted them with white noise. When 2^nd^-moment matrices are used as opposed to RSMs for characterizing the representational structures of the regions, the Riemannian distance gives larger effect sizes (z-scores) for all subjects.

#### Non-linear multivariate relationships

Next, we explored whether our suggested measure can also capture non-linear relationships. For that, we used Rectified Linear Units (ReLU) for the function *f*. The choice of nonlinearity was arbitrary here, but the results replicate for other types of nonlinear functions. Having said that, using ReLU was motivated by neural network models of brain information processing (Nair and Hinton, 2010). **Figure 15**(A-D) gives the simulation results for the different measures of representational connectivity. One can see that for the majority of parameter values, all measures give significant z-scores. However, when the ratio of the number of conditions to the number of voxels is small, dRiem gives higher z-scores. **Figure 15**E is color-coded based on the *winning measure(s)* for each combination of parameters (pooling across-metric tests for each combination of parameter values). We can clearly see that for relatively smaller ratios of the number of conditions to the number of voxels, dRiem outperforms all other measures, whereas when the number of conditions is large (e.g. when temporal geometries are being compared), the distance correlation performs best. In fact, in the range of relatively small condition to voxel count ratios all methods except for the Riemannian distance fail to even capture any connectivity. This is a unique advantage for the Riemannian distance when the designs are condition poor, or the ROIs consist of large numbers of voxels. For relatively larger number of conditions, where dCor performs best, other measures also give significant results. These are replicated for smaller number of simulated subjects (e.g. N = 20) as well.

**Figure 15.**
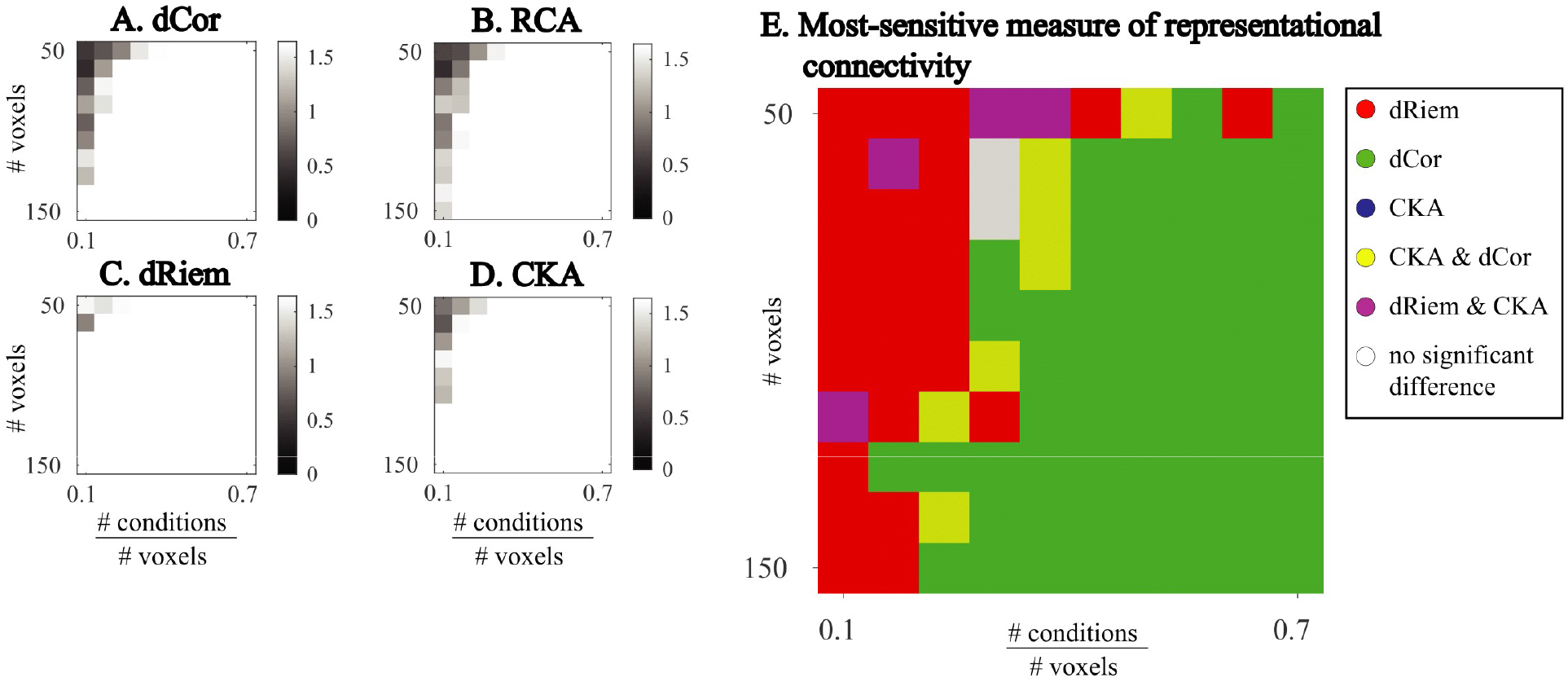
Comparing different measures of representational connectivity for simulated data with nonlinear interactions. *We simulated 50 subjects. For each subject a stimulus-response matrix in one region, X, was obtained by sampling from a multivariate normal distribution. The stimulus-response matrix in the other region, Y was ReLU*(*ReLU*(*X*) × *T*) *mimicking a typical nonlinear interaction found in neural network models. Each element of T was chosen randomly from a normal distribution. X and Y were then contaminated by i.i.d samples of additive Gaussian noise. Panels **A-D** represent the average z-scores of bias-corrected connectivity measures (across subjects). As one can see, in smaller condition to voxel count ratios, dRiem has larger z-scores, while in larger ratios, all metrics have z-scores that are equal to or larger than 1.645. This means that all of them are capable of detecting the nonlinear relationship. Panel **E** indicates the superiority regime for each metric. Pairwise comparisons between measures is done by performing an across-subject paired Wilcoxon signed-rank test on the two sets of z-scores (one per measure). Different measures are coded with different colors. If two measures are equally good at each voxel-condition count, their colors are combined (summed), and a new color is assigned (see the legend). For instance, magenta is the combination of blue (CKA), and red (dRiem) or white is the combination of all three colors. In this panel, no color is dedicated to RCA since it has the weakest performance among all metrics.*

### 3.6 The Riemannian distance outperforms existing measures in capturing similarity of neural network representations

One of the main advantages of RSA is that it can also be used to compare model representations, e.g., those in artificial neural networks (Kriegeskorte, 2015). In this section, we study the performance of the Riemannian distance in capturing similarity of neural network representations. Recently Kornblith and colleagues have proposed a measure called centered kernel alignment (CKA) for quantifying the similarity of neural network representations. Here, we introduce a similar approach and compare the Riemannian distance to CKA and other measures of representational similarity.

To this end, we perform almost the same analysis conducted in part 6.1 of (Kornblith et al., 2019). Consider two neural networks with a similar structure but different random initializations. The intuition underlying the method proposed by (Kornblith et al., 2019) is as follows: the stimulus-response matrix of a layer in the first network must have the highest similarity score to the stimulus-response matrix of the corresponding layer in the second network. For example, representations of layer 4 in network 1 must be maximally similar to representations of layer 4 in network 2, where networks 1 and 2 are trained with the same dataset but have different random initializations.

A VGG-like convolutional network based on All-CNN-C (Springenberg et al., 2015) is used for simulations. The detail of the architecture can be found in Table E.1 of (Kornblith et al., 2019). Since our purpose is to investigate the similarities of layer representations of neural networks trained with different random initializations, we trained the networks without extensive search for optimal hyperparameters. The average accuracy of ten networks in which the weights were initialized differently was acceptable (85.08%) when these networks were used on the test dataset.

We also used the CIFAR-10 dataset (Krizhevsky et al., 2009) which is composed of ten different categories of images. To construct the representations of different layers of the networks, we randomly sampled one image from each category from the training dataset. We used these ten images as input to different networks. The motivation for this was to resemble neuroscience experiments in which a random sample of all images is presented to the subject while brain responses are recorded. However, this is different from what (Kornblith et al., 2019) did in their work. They used all training and test data as inputs and compared representation for all those *conditions*.

For a given metric, in each pair of networks, if a layer of the first network has the most similar representation with its corresponding layer in the second network, we give a *score* of one to that layer and then calculate the average score for all of the layers. We repeat this procedure for all pairs of networks and calculate the average. Furthermore, we repeat the whole procedure with 100 different image samples and obtain the overall average. This is done to minimize the impact of the choice of the input images.

The results are depicted in **Figure 16**. The heights of the bars correspond to the average scores for each metric, and the error bars show the standard deviation of the scores across many choices of input images. As can be seen, in line with (Kornblith et al., 2019) findings, we also find that CKA performs better than RCA. However, the Riemannian distance has a significantly larger score than CKA and other measures. This shows that considering the geometry of the manifold of all 2^nd^-moment matrices can also have major contributions to neural network research.

**Figure 16.**
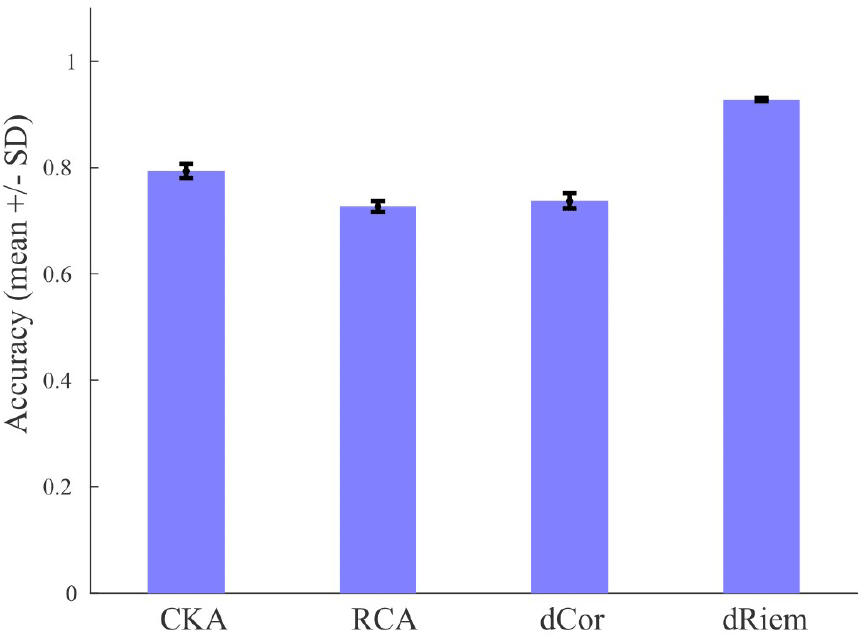
Accuracy of identifying corresponding layers based on maximum similarity in structurally similar 10-layer CNNs with different initializations. For each set of input images, which consists of one image from each category of images in the CIFAR-10 dataset (10 images in total), we obtain the score by examining each layer’s representational similarity to its corresponding layer in all pairs of networks. Each bar in this figure is the average of 100 scores obtained from 100 different choices of input images. The error bars are the standard deviations over many choices of input images.

### 3.7 The Riemannian metric can be successfully used for searchlight mapping with acceptable sensitivity

Having reported the advantages of the Riemannian framework for testing representational models in different brain regions, we next investigate its performance in continuous searchlight-based information mapping (Kriegeskorte et al., 2006).

One application of whole-brain searchlight analysis (Kriegeskorte et al., 2006) is to find brain regions in which RSMs are related to a reference RSM (e.g., a model or a reference RSM, *R*_*ref*_). In this analysis, for each subject, we compare *R*_*s*_, the RSM for a searchlight sphere, to *R*_*ref*_, and assign the normalized bias-corrected measure (i.e. the distance or correlation, divided by the standard deviation of the null distribution, see section 2.3 for details) between *R*_*s*_ and *R*_*ref*_ to the central voxel of the sphere. In our setup, matrices are 8 × 8 and we use twenty permutations^9^. This approach results in a normalized bias-corrected distance/correlation map per subject. We can then apply statistical tests to the extracted maps to test the summary statistics across subjects and obtain a p-map (e.g., via an across-subjects right-tailed signed-rank or t-test of bias-corrected values at each voxel). Finally, we use generalized Bonferroni correction (Neuwald and Green, 1994) to control the family-wise error rate at a threshold of 0.05 and select voxels from the p-map with p-values less than the corrected threshold as significant voxels.

Here, we use simulations (similar setup as (Nili et al., 2014)) in which a certain, restricted part of the brain contains a known multivariate effect (i.e., it has RSMs similar to a particular reference RSM). The green square in **Figure 17** illustrates a region with a known simulated effect. The voxels inside the green square have activity profiles coming from a specific known representational structure (RSM_in_) with some additive noise, while the activity profiles of other voxels do not follow any particular structure (RSM_out_).

**Figure 17.**
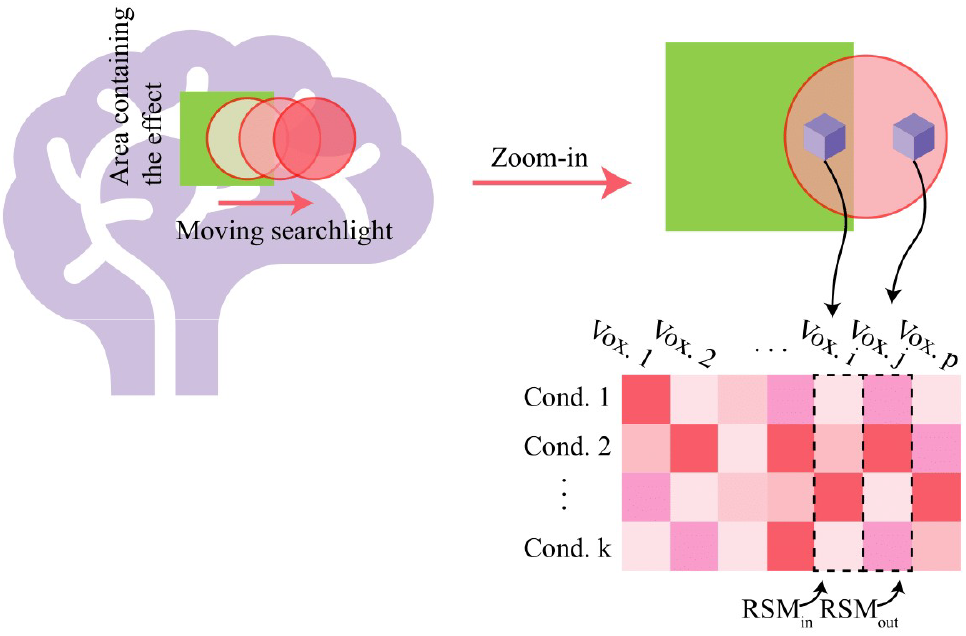
An illustration of the simulation approach for the searchlight-based analysis. The green square in the figure indicates an area of the brain with a specific representational structure: each voxel’s activity profile inside the green square follows RSM_in_. The responses of the other voxels consist of spatiotemporally smoothed Gaussian noise; therefore, their activity profiles and thus their RSM would be non-structured, RSM_out_. We then run a searchlight analysis with a model RSM that is identical to the one used for the green region. So, we expect to recover the green region after testing maps across subjects and also correcting for multiple comparisons (similar to any other whole-brain fMRI searchlight analysis).

The ideal metric would perfectly discriminate the green area from the rest. Therefore, after obtaining significant voxels from the method mentioned above, one can compare the performances of the metrics by their ability to detect the true underlying effect. We quantify the performance by the F1-score, which is widely accepted for quantifying accuracy for sparse effects (Sasaki, 2007). The score is the proportion of true positives over the sum of true positives and the average of false positives and false negatives.

**Figure 18** shows the output for the Riemannian metric when 20 subjects are simulated. One can observe that the analysis does, indeed, retrieve the simulated effect. It is worth noting that, to the best of our knowledge, this is the first attempt to test representational models with distance-based measures (like the Riemannian distance). The results reassure that the bias-correction step that we apply is valid and has good sensitivity and specificity.

**Figure 18.**
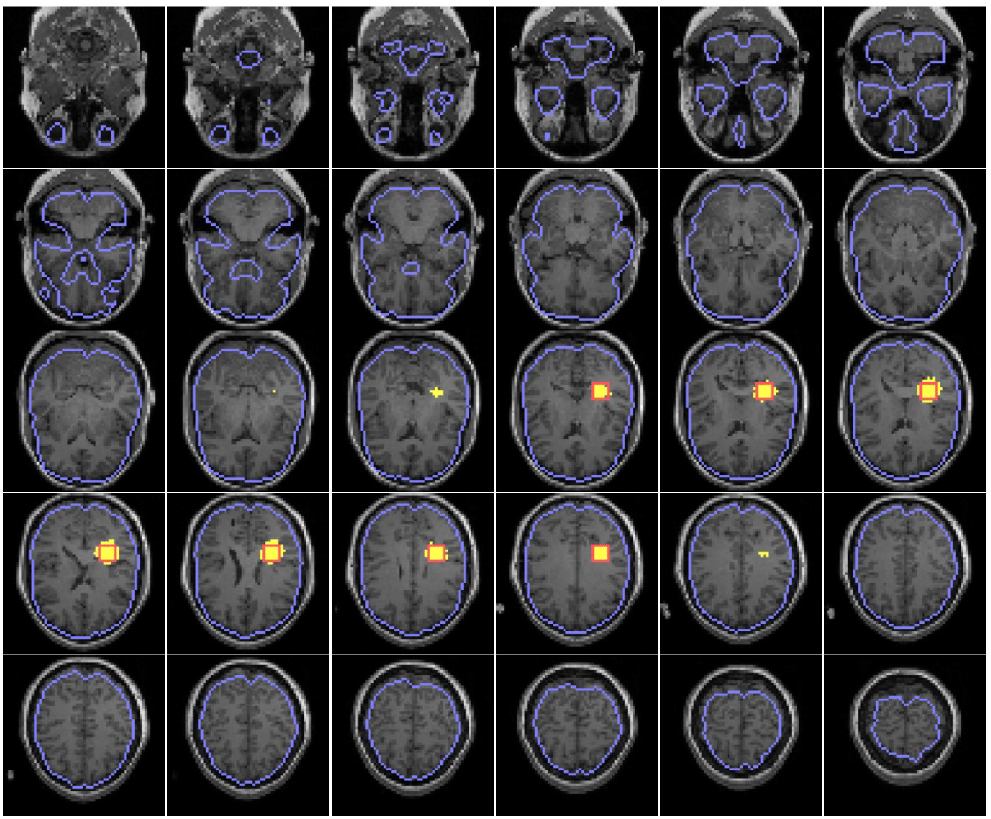
Sample prediction of related voxels using the Riemannian metric. Twenty subjects are simulated, and related voxels are detected using the Riemannian metric. Yellow voxels show the prediction, and red lines are the borders for the cubic ground truth effect. The family-wise error rate is controlled at 0.05.

In **Figure 19**, we have quantitatively compared the performances of different metrics when the number of simulated subjects is 20. Here we have simulated eight independent groups of subjects, each containing 20 subjects. For each group, we have obtained a p-map according to the procedure explained above and kept the significance threshold at a constant level of 0.05 after correcting for multiple testing. The boxplot shows how the F1-score of detected significant voxels varies among different simulated groups for each metric. One can see that the Riemannian metric shows superior performance to the Frobenius norm of difference and the Pearson correlation. The superiority over the Frobenius norm is another piece of evidence that shows considering the manifold of RSMs is important. Further, it has comparable performance to state-of-the-art metrics such as the Spearman or Kendall’s *τ* correlation. Here the superiority is tested by the paired right-tailed signed-rank test of F1-scores of two metrics calculated in different groups and shown by horizontal lines on top of the figure. It should be noted that we have used normalized bias-corrected values for all metrics in our analysis in order to have a fair comparison; however, it is possible to use raw values for correlation-based metrics without losing performance (this implicitly assumes that correlations are zero-centered under the null hypothesis).

**Figure 19.**
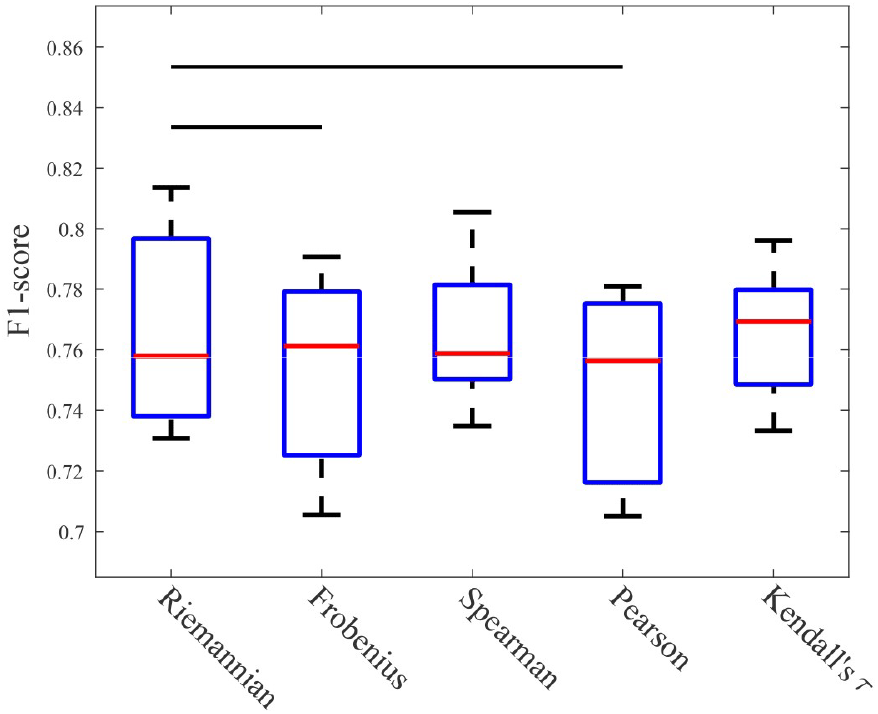
Comparison of F1-scores for different metrics. For 20 subjects and a metric, a p-map is obtained by testing normalized bias-corrected maps across subjects. Then, voxels with a p-value less than corrected thresholds for the significance level of 0.05 are labeled as related (significant) voxels. The F1-score of predictions are shown for each metric. Here, we have repeated the procedure for eight independent groups of subjects, each containing 20 subjects. The proposed Riemannian metric shows comparable performance to the commonly used correlation-based methods (Spearman and Kendall’s τ) and is superior to others. Here, each horizontal bar shows the F1-scores obtained from different groups with the Riemannian metric are significantly larger than the other metrics when a right-tailed signed-rank test is performed.

Taken together, these results suggest that the Riemannian distance can be used successfully for searchlight mapping. Since distances are positively biased, inference on the Riemannian distance requires correcting for the bias at the single-subject level, and that increases the computational load of the analysis. Future work is needed to allow accurate and computationally simple estimation of the null distribution.

## 4 Discussion and Conclusions

Pattern information analysis has played an important role in understanding the information content of brain representations and gaining insight into the underlying neural computations (Haynes and Rees, 2006; Kriegeskorte and Diedrichsen, 2019; Quian Quiroga and Panzeri, 2009). Apart from pattern classifiers, other representational methods rely on the 2^nd^-moment matrix of stimulus-evoked activity patterns (Diedrichsen and Kriegeskorte, 2017). Representational dissimilarity matrices (RDMs) and stimulus-by-stimulus correlation matrices of activity patterns (RSMs) can both be directly derived from 2^nd^-moment matrices.

RSA has been widely used in studies that test hypotheses about brain representations. However, inference procedures to date have been blind to the manifold that underlies the 2^nd^-moment matrices, RSMs, and positive semi-definite (PSD) matrices in general. The Riemannian manifold has been used in other fields of neuroscience research to better capture the relationship between PSD matrices. As both 2^nd^-moment matrices and RSMs also reside on the Riemannian manifold, we investigate whether the framework can offer improvements to the classic RSA inference (Kriegeskorte et al., 2008; Nili et al., 2014).

Since the Euclidean and Riemannian spaces employ different metrics, the relationship between sets of activity patterns, from brains or models, can be different depending on the metric. For example, we illustrated a case (section 1) where the similarity of representations is completely different in the Euclidean space and the Riemannian manifold.

Given this difference, we compared distances on the Riemannian manifold with other existing metrics and found many advantages for adopting the Riemannian geometry into the RSA.

First, one can think of activity profiles as points (e.g., corresponding to response channels in brain measurements) in a space where each axis corresponds to one experimental condition (e.g. stimulus). The more samples one can get from a brain region, the more accurate the estimates of the variance of the points or the 2^nd^-moment matrix will be. In fact, with inadequate sampling, two regions that have different functional properties can have identical fits to the same model. The discrepancy between the two regions becomes more and more clear with increasing samples. We showed that the Riemannian distance is less susceptible to this sampling bias.

Second, we explored the extent to which Riemannian distances are smaller for measurements of the same set of stimuli for a participant (within-subject reliability). We followed the same line of reasoning as (Walther et al., 2016), but instead of comparing different estimates of pattern dissimilarity, compared different RDM similarity measures. Also, we performed the comparison for two different datasets. In one, we used anatomical ROIs, and the other, we applied the analysis to functional ROIs (from an independent localizer run). We observed that in both datasets, comparisons made on the basis of the Riemannian distance yield significantly higher split-data reliability across different ROIs. Next, we provided further evidence on the superiority of the Riemannian distance by showing that it allows discriminating between representational matrices of disjoint areas of the brain as well.

It might be that in these cases, different representational matrices of a subject in a brain region become farther apart in the Euclidean space but less in the Riemannian manifold. Future work can characterize the curvature of the Riemannian manifold and identify regions of the manifold in which relationships vary considerably depending on the space (i.e., localize parts of the manifold where Euclidean relationships would be maximally different from relationships on the Riemannian manifold).

Making inferences on distance-based metrics faces a major difficulty. While with RDM correlations, it can be assumed that the null values are centered around zero, this type of assumption does not apply to distances. Distances are always positive, and this complicates statistical inferences on their values. Here, we followed and extended the approach taken by (Basti et al., 2020) and showed that it is indeed possible to test for significant similarities if each measure is *normalized* by its null value. This *bias correction* scheme allows for making inference on measures that have different scales and different null distributions. Our preliminary investigation shows that with 20 subjects, the results do not change considerably if the number of permutations goes as low as 20. However, this might vary with the number of subjects and needs further work to become part of the RSA inference pipeline.

Brain regions do not function in isolation, and connectivity methods quantify *communication* between brain regions. Multi-dimensional connectivity methods test for connectivity in the information carried by different multi-dimensional measurements (Basti et al., 2020). That is in contrast to the classic univariate functional connectivity that tests the correlation between average time series of two regions (Biswal et al., 1995). Given the benefits of using the Riemannian metric that we observed for model testing, we next considered its applications in connectivity analysis. We first validated our metric by showing that it can capture multivariate linear relationships with high accuracy. Then, we showed that the metric can be particularly useful for testing nonlinear regional interactions. Once again, the intuition is that nonlinear relationships can considerably move RSMs or 2^nd^-moment matrices in the Euclidean space, but not so much when the underlying Riemannian manifold is considered. Additionally, we showed that testing the relationship between 2^nd^-moment matrices, as opposed to RSMs, is more powerful in this context. 2^nd^-moment matrices also contain information about regional average activations (univariate effects), and therefore, our proposed measure of functional connectivity incorporates both univariate and multivariate connectivity.

Another motivation for using the Riemannian distance in functional connectivity is its affine-invariant property (section 3.4). Relationships between 2^nd^-moment matrices are unaffected by identical full-rank linear transformations. This property can be useful for any case where sources are combined to obtain measured signals (Pham & Cardoso, 2001).

One might note that a caveat of our proposed metric is that it requires the number of conditions to be less than the number of voxels. Future work can develop ways to afford analysis on the Riemannian manifold for the cases in which the number of conditions are larger than the number of voxels (surely, one way to approach that would be to reduce the dimensionality of the data, though this might not be ideal in many scenarios).

Researchers are often interested in localizing multivariate effects in locally distributed activity patterns. Searchlight mapping is best suited for this purpose. Using simulations with known ground truth, we verified that the Riemannian metric works well in searchlight analysis. We showed that while it has an acceptable false positive rate, its sensitivity is either higher or equal to existing methods. Given the advantages the we have reported for the Riemannian metric, it is completely plausible that real data applications of searchlight analysis would find cases in which only the Riemannian metric can detect the presence of an effect. For example, in our simulations, response patterns in a seed region were linearly related to a particular structure. However, non-linear relationships will be best captured by the Riemannian metric (section 3.5). Thus, although our searchlight simulations were not designed so that the benefits of the Riemannian metric are best highlighted, we still find that the metric performs better and, at a minimum yields results equal to the existing methods.

The analyses performed in this manuscript are conducted for both RSMs and 2^nd^-moment matrices (G) when possible. 2^nd^-moment matrices are more general and summarize the information present in response patterns as well as in regional average activations (diagonal elements of G). Similar to PCM (Diedrichsen et al., 2011), model testing in RSA using 2^nd^-moment matrices can lead to testing a hypothesis about functional properties that are both sensitive to univariate and multivariate distinctions. It is also plausible that model testing with G can allow the separation of the contributions of univariate average responses and fine-grained pattern effects when used in discriminating different brain states. This, in itself, can help in understanding the neural code. Future work will consolidate this and apply it to empirical data.

One main advantage of RSA is that testing computational models is integrated with the analysis of neuroimaging data. An example of this would be a testing hypothesis about RDMs/RSMs from different layers of neural networks (Khaligh-Razavi and Kriegeskorte, 2014) and how they explain brain functions (Kriegeskorte, 2015). We reasoned that if the Riemannian framework has a lot to offer to test brain data, it might also be useful in capturing the relationship between representations of artificial neural networks. We verified that the Riemannian metric is best suited for capturing the similarity of model representations. In particular, we showed that it performs significantly better when compared to a recent novel metric that was shown to be most useful in capturing representational similarity (Kornblith et al., 2019).

We believe that the results from our neural network analysis go hand in hand with the other results we report in this paper. Using human data, we showed that different samples of each brain region of a subject are more similar to each other and more distinct from other regions of the same subject when the Riemannian distance is used to compare representations. With the deep-nets, model features from the same layer of networks with different initializations could be thought of as similar to brain responses of an ROI for different subjects. Future work can verify whether different initializations can explain individual differences, but in our results this intuition can link results from real data and neural network analysis (section 3.6).

Although future work is required to pinpoint a priori conditions in which the Riemannian distance outperforms other metrics, we showed that the Riemannian geometry could be extremely helpful for pattern information analysis. We believe that the Riemannian geometry has a lot to offer to multivariate pattern analysis and in particular, RSA.

## Data and code availability

Links of the datasets we used in this study:

Visual object recognition dataset: https://openneuro.org/datasets/ds000105/versions/00001
Generic object decoding dataset: https://openneuro.org/datasets/ds001246/versions/1.2.1

Analyses scripts are publicly available at https://github.com/mshahbazi1997/riemRSA

## Footnotes

1 A stimulus-by-stimulus correlation matrix, also sometimes referred to as a representational similarity matrix (RSM), is 1 - correlation-distance RDM.

2 This requires assuming that the origin is the same as the centre of the data points in the multivariate response space.

3 *trace*(*AB*) = *trace*(*BA*)

4 All *k* × *k* PSD matrices lie on the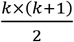-dimensional surface. 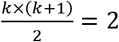 does not have an answer for *k* ∈ ℕ.

5 Consider a region with a certain volume. Choosing different voxel sizes would result in a different number of voxels for the ROI.

6 This was part of the motivation for defining the metric (see section 2.2 for the mathematical derivation).

7 Alternatively, we could have also computed the Riemannian distance between their corresponding RSMs. This distinction is elaborated later in this section.

8 In the previous sections we intentionally included RSMs to relate our measure to classic RSA studies.

9 Larger numbers of permutations had little effect on improving results.

## References

Allefeld, C., Görgen, K., Haynes, J.D., 2016. Valid population inference for information-based imaging: From the second-level t-test to prevalence inference. Neuroimage. https://doi.org/10.1016/j.neuroimage.2016.07.040

Barachant, A., Bonnet, S., Congedo, M., Jutten, C., 2012. Multiclass brain-computer interface classification by Riemannian geometry. IEEE Trans. Biomed. Eng. https://doi.org/10.1109/TBME.2011.2172210

Basti, A., Nili, H., Hauk, O., Marzetti, L., Henson, R.N., 2020. Multi-dimensional connectivity: a conceptual and mathematical review. Neuroimage 221, 117179. https://doi.org/10.1016/j.neuroimage.2020.117179

Benjamini, Y., Hochberg, Y., 1995. Controlling the False Discovery Rate: A Practical and Powerful Approach to Multiple Testing. J. R. Stat. Soc. Ser. B. https://doi.org/10.1111/j.2517-6161.1995.tb02031.x

Bhatia, R., 2009. Positive definite matrices, Positive Definite Matrices. https://doi.org/10.2307/2317709

Biswal, B., Zerrin Yetkin, F., Haughton, V.M., Hyde, J.S., 1995. Functional connectivity in the motor cortex of resting human brain using echo-planar mri. Magn. Reson. Med. https://doi.org/10.1002/mrm.1910340409

Carmo, M.P. do, 1992. Riemannian Geometry, Riemannian Geometry. https://doi.org/10.1007/978-1-4757-2201-7

Congedo, M., Barachant, A., Bhatia, R., 2017. Riemannian geometry for EEG-based brain-computer interfaces; a primer and a review. Brain-Computer Interfaces. https://doi.org/10.1080/2326263X.2017.1297192

Coutanche, M.N., Thompson-Schil, S.L., 2013. Informational connectivity: Identifying synchronized discriminability of multi-voxel patterns across the brain. Front. Hum. Neurosci. https://doi.org/10.3389/fnhum.2013.00015

Diedrichsen, J., Kriegeskorte, N., 2017. Representational models: A common framework for understanding encoding, pattern-component, and representational-similarity analysis. PLoS Comput. Biol. https://doi.org/10.1371/journal.pcbi.1005508

Diedrichsen, J., Ridgway, G.R., Friston, K.J., Wiestler, T., 2011. Comparing the similarity and spatial structure of neural representations: A pattern-component model. Neuroimage. https://doi.org/10.1016/j.neuroimage.2011.01.044

Gretton, A., Bousquet, O., Smola, A., Scḧlkopf, B., 2005. Measuring statistical dependence with Hilbert-Schmidt norms, in: Lecture Notes in Computer Science (Including Subseries Lecture Notes in Artificial Intelligence and Lecture Notes in Bioinformatics). https://doi.org/10.1007/11564089_7

Hanson, S.J., Matsuka, T., Haxby, J. V., 2004. Combinatorial codes in ventral temporal lobe for object recognition: Haxby (2001) revisited: Is there a “face” area? Neuroimage. https://doi.org/10.1016/j.neuroimage.2004.05.020

Haxby, J. V., Gobbini, M.I., Furey, M.L., Ishai, A., Schouten, J.L., Pietrini, P., 2001. Distributed and overlapping representations of faces and objects in ventral temporal cortex. Science (80-.). https://doi.org/10.1126/science.1063736

Haynes, J.D., Rees, G., 2006. Decoding mental states from brain activity in humans. Nat. Rev. Neurosci. https://doi.org/10.1038/nrn1931

Horikawa, T., Kamitani, Y., 2017. Generic decoding of seen and imagined objects using hierarchical visual features. Nat. Commun. https://doi.org/10.1038/ncomms15037

Kay, K.N., Naselaris, T., Prenger, R.J., Gallant, J.L., 2008. Identifying natural images from human brain activity. Nature. https://doi.org/10.1038/nature06713

Khaligh-Razavi, S.M., Kriegeskorte, N., 2014. Deep Supervised, but Not Unsupervised, Models May Explain IT Cortical Representation. PLoS Comput. Biol. https://doi.org/10.1371/journal.pcbi.1003915

Kornblith, S., Norouzi, M., Lee, H., Hinton, G., 2019. Similarity of neural network representations revisited, in: 36th International Conference on Machine Learning, ICML 2019.

Kriegeskorte, N., 2015. Deep Neural Networks: A New Framework for Modeling Biological Vision and Brain Information Processing. Annu. Rev. Vis. Sci. https://doi.org/10.1146/annurev-vision-082114-035447

Kriegeskorte, N., Diedrichsen, J., 2019. Peeling the Onion of Brain Representations. Annu. Rev. Neurosci. https://doi.org/10.1146/annurev-neuro-080317-061906

Kriegeskorte, N., Goebel, R., Bandettini, P., 2006. Information-based functional brain mapping. Proc. Natl. Acad. Sci. U. S. A. https://doi.org/10.1073/pnas.0600244103

Kriegeskorte, N., Kievit, R.A., 2013. Representational geometry: Integrating cognition, computation, and the brain. Trends Cogn. Sci. https://doi.org/10.1016/j.tics.2013.06.007

Kriegeskorte, N., Mur, M., Bandettini, P., 2008. Representational similarity analysis - connecting the branches of systems neuroscience. Front. Syst. Neurosci. https://doi.org/10.3389/neuro.06.004.2008

Krizhevsky, A., Nair, V., Hinton, G., 2009. CIFAR-10 and CIFAR-100 datasets [WWW Document]. https://www.cs.toronto.edu/~kriz/cifar.html.

Nair, V., Hinton, G.E., 2010. Rectified linear units improve Restricted Boltzmann machines, in: ICML 2010 - Proceedings, 27th International Conference on Machine Learning.

Neuwald, A.F., Green, P., 1994. Detecting patterns in protein sequences. J. Mol. Biol. 239, 698–712.

Nichols, T.E., Holmes, A.P., 2002. Nonparametric permutation tests for functional neuroimaging: A primer with examples. Hum. Brain Mapp. https://doi.org/10.1002/hbm.1058

Nili, H., Walther, A., Alink, A., Kriegeskorte, N., 2020. Inferring exemplar discriminability in brain representations. PLoS One. https://doi.org/10.1371/journal.pone.0232551

Nili, H., Wingfield, C., Walther, A., Su, L., Marslen-Wilson, W., Kriegeskorte, N., 2014. A Toolbox for Representational Similarity Analysis. PLoS Comput. Biol. 10, e1003553. https://doi.org/10.1371/journal.pcbi.1003553

O’Toole, A.J., Jiang, F., Abdi, H., Haxby, J. V., 2005. Partially distributed representations of objects and faces in ventral temporal cortex. J. Cogn. Neurosci. https://doi.org/10.1162/0898929053467550

Panzeri, S., Treves, A., 1996. Analytical estimates of limited sampling biases in different information measures. Netw. Comput. Neural Syst. https://doi.org/10.1088/0954-898X/7/1/006

Pennec, X., 2006. Intrinsic statistics on Riemannian manifolds: Basic tools for geometric measurements. J. Math. Imaging Vis. https://doi.org/10.1007/s10851-006-6228-4

Pennec, X., Fillard, P., Ayache, N., 2006. A riemannian framework for tensor computing. Int. J. Comput. Vis. https://doi.org/10.1007/s11263-005-3222-z

Pennec, X., Sommer, S., Fletcher, T., 2019. Riemannian Geometric Statistics in Medical Image Analysis, Riemannian Geometric Statistics in Medical Image Analysis. https://doi.org/10.1016/C2017-0-01561-6

Pervaiz, U., Vidaurre, D., Woolrich, M.W., Smith, S.M., 2020. Optimising network modelling methods for fMRI. Neuroimage. https://doi.org/10.1016/j.neuroimage.2020.116604

Pham, D.T., Cardoso, J.F., 2001. Blind separation of instantaneous mixtures of nonstationary sources. IEEE Trans. Signal Process. https://doi.org/10.1109/78.942614

Quian Quiroga, R., Panzeri, S., 2009. Extracting information from neuronal populations: Information theory and decoding approaches. Nat. Rev. Neurosci. https://doi.org/10.1038/nrn2578

Rahim, M., Thirion, B., Varoquaux, G., 2019. Population shrinkage of covariance (PoSCE) for better individual brain functional-connectivity estimation. Med. Image Anal. https://doi.org/10.1016/j.media.2019.03.001

Sasaki, Y., 2007. The truth of the F-measure. Teach Tutor mater 1–5.

Springenberg, J.T., Dosovitskiy, A., Brox, T., Riedmiller, M., 2015. Striving for simplicity: The all convolutional net. 3rd Int. Conf. Learn. Represent. ICLR 2015 - Work. Track Proc. 1–14.

Székely, G.J., Rizzo, M.L., 2014. Partial distance correlation with methods for dissimilarities. Ann. Stat. 42, 2382–2412. https://doi.org/10.1214/14-AOS1255

Walther, A., Nili, H., Ejaz, N., Alink, A., Kriegeskorte, N., Diedrichsen, J., 2016. Reliability of dissimilarity measures for multi-voxel pattern analysis. Neuroimage. https://doi.org/10.1016/j.neuroimage.2015.12.012

You, K., Park, H.J., 2021. Re-visiting Riemannian geometry of symmetric positive definite matrices for the analysis of functional connectivity. Neuroimage. https://doi.org/10.1016/j.neuroimage.2020.117464

